# The Functional Logic of Odor Information Processing in the *Drosophila* Antennal Lobe

**DOI:** 10.1101/2021.12.27.474306

**Authors:** Aurel A. Lazar, Tingkai Liu, Chung-Heng Yeh

**Author notes:** The authors’ names are listed in alphabetical order.

## Abstract

The early olfactory system of the fruit fly, while sensing a complex odorant landscape, encodes the *odorant object identity* (semantic information) and *the odorant concentration waveform* (syntactic information) into a combinatorial neural code. Single-channel physiology recordings at the output of the Antenna Lobe (AL) exhibit *concentration-invariance* and *contrast-boosting* properties, indicating a decoupling of the odorant object identity from the concentration waveform in steady-state while responding strongly to odorant concentration onset and offset in transient states.

Through exhaustive computational explorations of the AL circuits, we show that the steady-state and transient response features of the AL are, respectively, due to presynaptic and postsynaptic Local Neurons (LNs). Theoretical analysis reveals that the LN pathways can be modeled as parallel differential Divisive Normalization Processors (DNPs). Differential DNPs robustly extract odorant identity (semantic information) and ON/OFF odorant event-timing (syntactic information), thereby providing for the AL the functional logic of *ON-OFF odorant identity recovery*.

## 1 Introduction

The early olfactory system of the fruit fly senses a complex odorant landscape [1], encoding both the *odorant object identity* and *the odorant concentration waveform* [2] into a combinatorial neural code at the level of the axons of the Olfactory Sensory Neurons (OSNs) of the Antenna. Odor information processing in the Antennal Lobe (AL) and its resulting representation by the Projection Neurons (PNs), in turn, is further processed in higher brain centers for recognition, associative learning, and other cognitive tasks [3, 4, 5].

Odorant object identity is defined by two parameters [2]:

1. the *binding rate* [**b**]*_ron_*, i.e., the strength a given odorant molecule *o* binds to the receptors of type *r* = {1, 2,…, *R*} expressed by OSN *n*, and
2. the *dissociation rate* [**d**]*_ron_*, i.e., the rate at which the odorant molecule *o* dissociates from its previously bound odorant receptors of type *r* expressed by OSN *n*.

As olfactory stimuli are transmitted via odorant plumes with a time varying concentration profile, the odorant sensory input is encoded by multiplicative coupling between the odorant object identity and the concentration amplitude waveform *u*(*t*) [6, 2] (see **Fig.**1(A)):

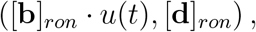

which in turn is transduced into a spike train by the OTP-BSG cascade of the OSN [2]. The multiplicative coupling underlying the encoding process results in a confounding OSN representation of identity and concentration [7, 8]. Consequently, robust odorant recognition and olfactory associative learning calls for the AL circuit to largely generate concentration-invariant PN spatio-temporal responses across odorant object identities (**Fig.**1(A)) [9, 10, 11].

**Figure 1:**
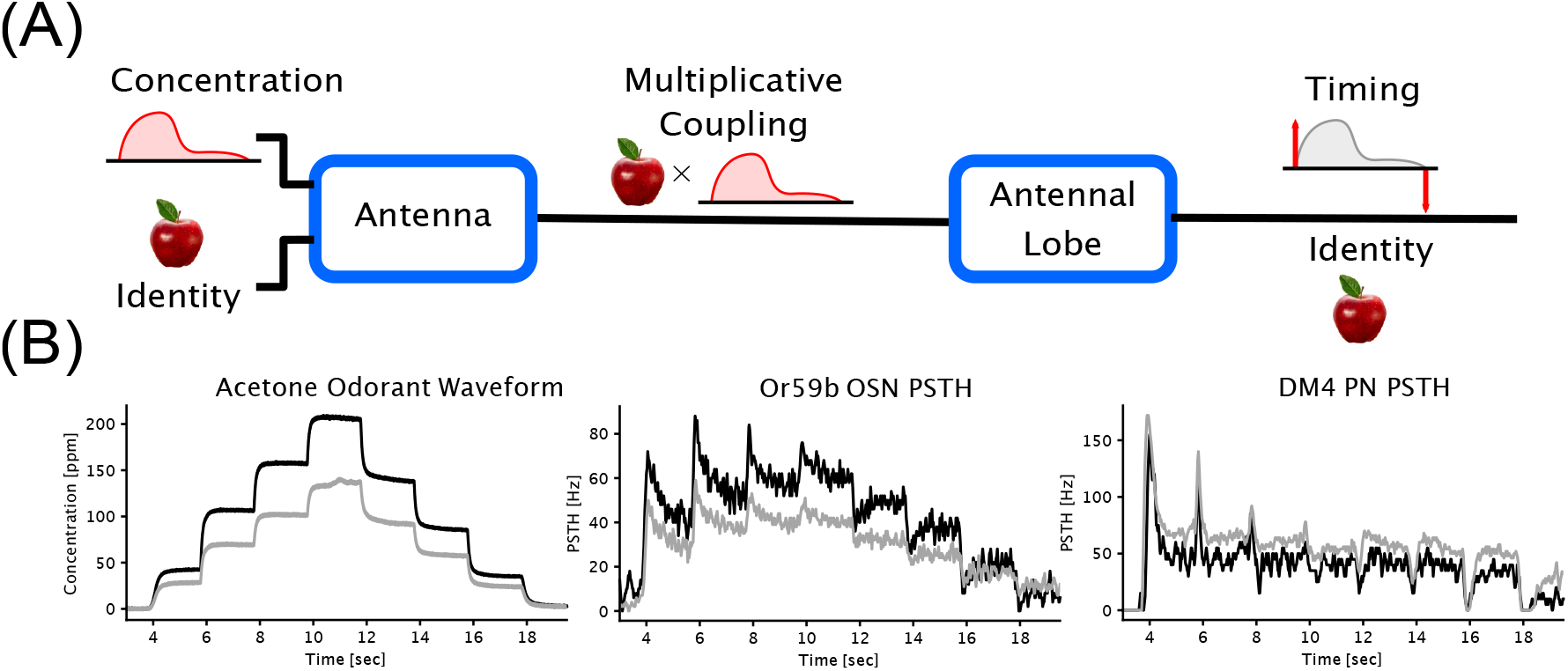
Odor signal processing in the Antennal Lobe circuit of the fruit fly brain. (A) Computational logic of the olfactory pathway. Encoding of odorant waveforms in the Antenna is confounded between odorant identity and concentration. For robust downstream odor signal processing, recognition and associative learning, the Antennal Lobe transforms the confounded Antenna odorant representation into on/offset timing information and odorant identity recovery. (B) Input/Output (I/O) characteristics of the early olfactory pathway. Shown from left to right are Acetone staircase odorant waveforms, Or59b OSN PSTHs and DM4 PN PSTHs, respectively. Data taken from [23].

Structurally, the axons of OSNs expressing the same receptor type [12, 6] and the dendrites of downstream PNs form ~ 50 glomeruli [13, 14, 12], a set of parallel *channels* of odor signal processing organized by receptor types [3]. An example input/output map along a single channel is shown in **Fig.**1(B). Importantly, computations both within and between channels in the Antennal Lobe are facilitated by about 120 local neurons (LNs) [15, 16, 17, 18, 19, 20] that are differentiated by their innervation patterns (within vs. across glomeruli), neurotransmitter types, excitatory/inhibitory synapses, and innervation targets. Distilling connectivity patterns from the connectomes of both the adult [21] and larva [22] *Drosophila*, we considered 3 main LN cell types: 1) pre-synaptic pan-glomerular (innervating all glomeruli) inhibitory LNs (Pre-LNs), 2) post-synaptic uni-glomerular (innervating a single glomerulus) excitatory LNs (Post-eLNs), and 3) post-synaptic uni-glomerular inhibitory LNs (Post-iLNs).

Thus, the challenges arise as to what are the nature of 1) the computational role played by the LNs in generating the PN physiological responses, and 2) the functional logic underlying such computations.

Addressing the challenges with a computational approach, we exhaustively evaluated single-channel and multi-channel AL circuit models both across variations of LN connectivity (local vs. global, feedforward vs. feedback) and across random samplings of model parameters for each circuit.

To comparatively analyze the circuits and the computational role of each LN type, we first note that decomposition of the OSN and PN responses into steady-state and transient components (**Fig.**2(B)) revealed important differences in response features. In steady-state, single-channel PN responses exhibit a significantly reduced dependency of the amplitude of the concentration waveform (see also **Fig.**11(A bottom)), phenomenon henceforth referred to as *concentration-invariance* [9, 10, 11, 8, 7]. Extending such phenomenon to multi-channel AL circuits, we hypothesize that steady-state population PN responses recover the *odorant object identity* as represented by the odorant *affinity rate* [**b**]*_ron_*/[**d**]*_ron_*. On the other hand, transient PN responses show a much higher correlation with the concentration contrast (**Fig.**2(C) and **Fig.**11(B middle), see Methods for definition of concentration contrast), a phenomenon henceforth referred to as *contrast-boosting*.

**Figure 2:**
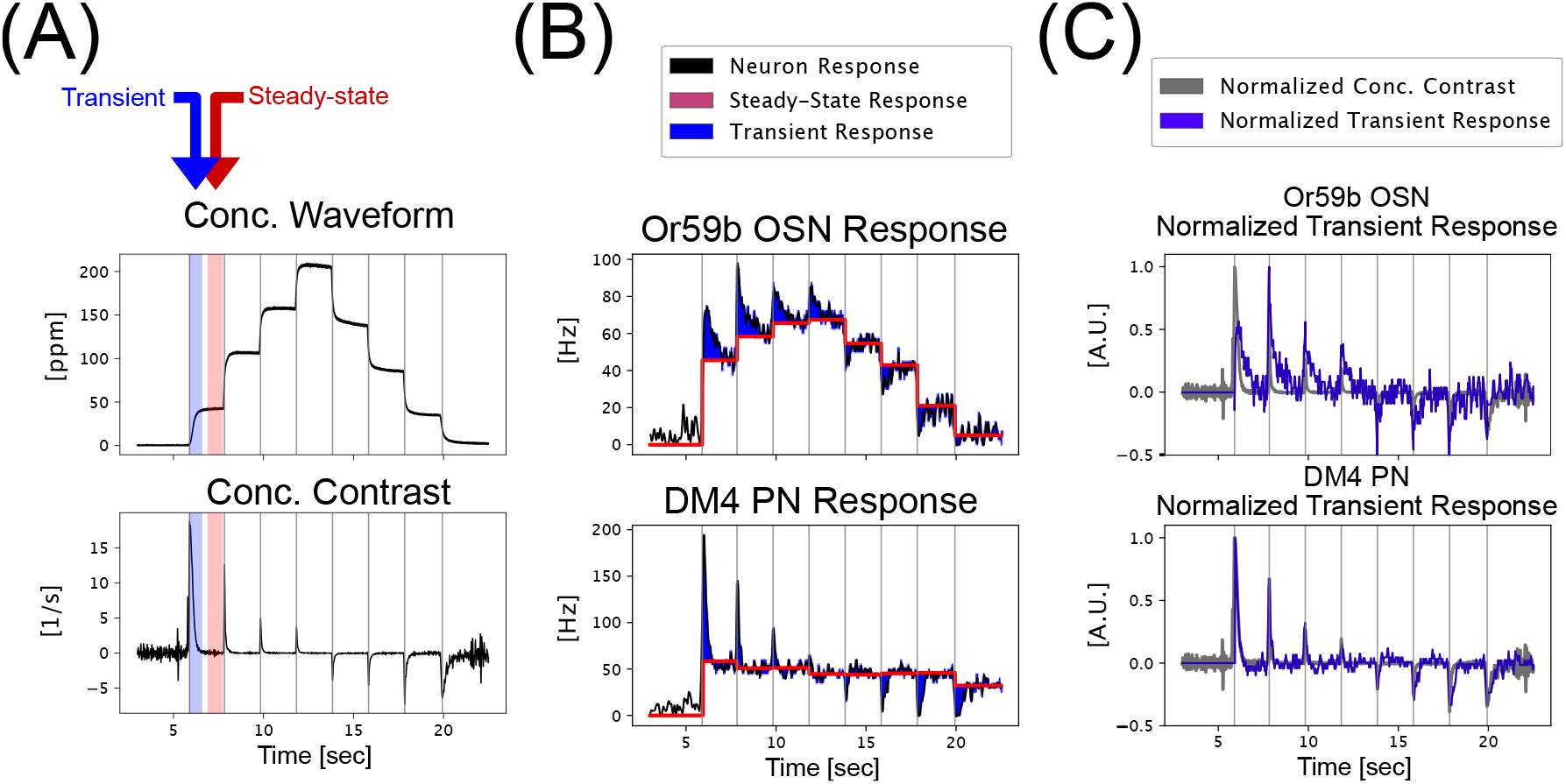
Essential temporal encoding/processing characteristics of the DM4 PNs. (A) Acetone concentration waveform (first published in [23]) and concentration contrast. (B) Or59b OSN and DM4 PN spike train data in response to an Acetone staircase waveform. Each neuron response is decomposed into steady-state and transient responses - where the steady-state response is computed as the average PSTH within a 500 miliseconds window before a jump in odorant concentration. The transient response is the residual obtained by subtracting the steady-state response from the overall response. See the text for more details. (C) Neuron normalized transient response compared against normalized concentration contrast (bottom). Note that the PN transient response agrees very well with the concentration contrast.

Using these PN response characteristics as metrics, we found that presynaptic (to OSN-to-PN synapse) LN inhibition across glomeruli is essential for enhancing the *concentration-invariance* of the PN responses, thereby stably recovering odorant object identities as odorant affinity vectors. On the other hand, postsynaptic (to OSN-to-PN synapse) LN excitation and inhibition capture odorant concentration waveform onset and offset timing information, strongly boosting the contrast of the PN transient responses from the ones observed in the OSNs responses.

Addressing the challenges with a theoretical approach, we demonstrate that the AL circuit can be described by a functionally equivalent circuit with 3 parallel differential Divisive Normalization Processors (DNPs) [24], extracting the concentration-invariance and ON/OFF contrast-boosting features independently of each other. From a functional perspective, such separation of identity and concentration information suggests an *ON-OFF odorant object identity recovery* paradigm for the AL circuit architecture. Such a paradigm enables rapid and robust odorant recognition and olfactory associative learning downstream.

## 2 Results

To discover the functional logic of the Antennal Lobe, we employed comparative analysis of temporal **Section** 2.1 and spatio-temporal **Section** 2.2 AL circuits with different LN configurations. In **Section** 2.3, we show that the three LN pathways (presynaptic inhibition, postsynaptic inhibition & excitation) are computationally equivalent to three differential Divisive Normalization Processors (DNPs). Collectively, the three DNPs support an *ON-OFF odorant object identity recovery* processing paradigm in the Antennal Lobe that is robust to different odorant objects and concentration waveforms (**Section** 2.4).

### 2.1 Pre-/Post-synaptic Local Neurons Control Concentration Invariance/Contrast Boosting of Single Channels

Recall that analysis of the temporal single channel PN response reveals two key features of odor signal processing: *concentration invariance* and *contrast boosting*. To systematically determine the neural basis of these processing features implemented by Pre-LNs, Post-eLNs and Post-iLNs, we exhaustively compared all architectural variations of the AL circuit. In particular, we compared, on the pre-synaptic side of the OSN-to-PN synapse, three different configurations of Pre-LN inhibition of the OSN Axon-Terminal: 1) no Pre-LNs (Baseline Circuit, **Fig**.3(A[i])), 2) Pre-LNs receiving input from the OSN Axon-Terminal and inhibiting, in turn, the OSN Axon-Terminal (Local Feedback Circuit, **Fig.**3(A[ii])), and 3) Pre-LNs receiving input from the OSN Axon-Hillock and inhibiting the OSN Axon-Terminal (Local Feedforward Circuit, **Fig**.3(A[iii])). On the post-synaptic side of the OSN-to-PN synapse, we consider three configurations of Post-eLNs and Post-iLNs: 1) no Post-eLNs or Post-iLNs (Baseline Circuit, **Fig.**3(A[i])), 2) only Post-eLNs (**Fig.**3(A[iv])), 3) only Post-iLNs(**Fig**.3(A[v])), 4) both Post-eLNs and Post-iLNs (**Fig**.3(A[vi])). Combining both pre-synaptic and post-synaptic configurations, a total of 12 circuit architectures were considered, 8 of which are shown in **Fig.**3(A).

**Figure 3:**
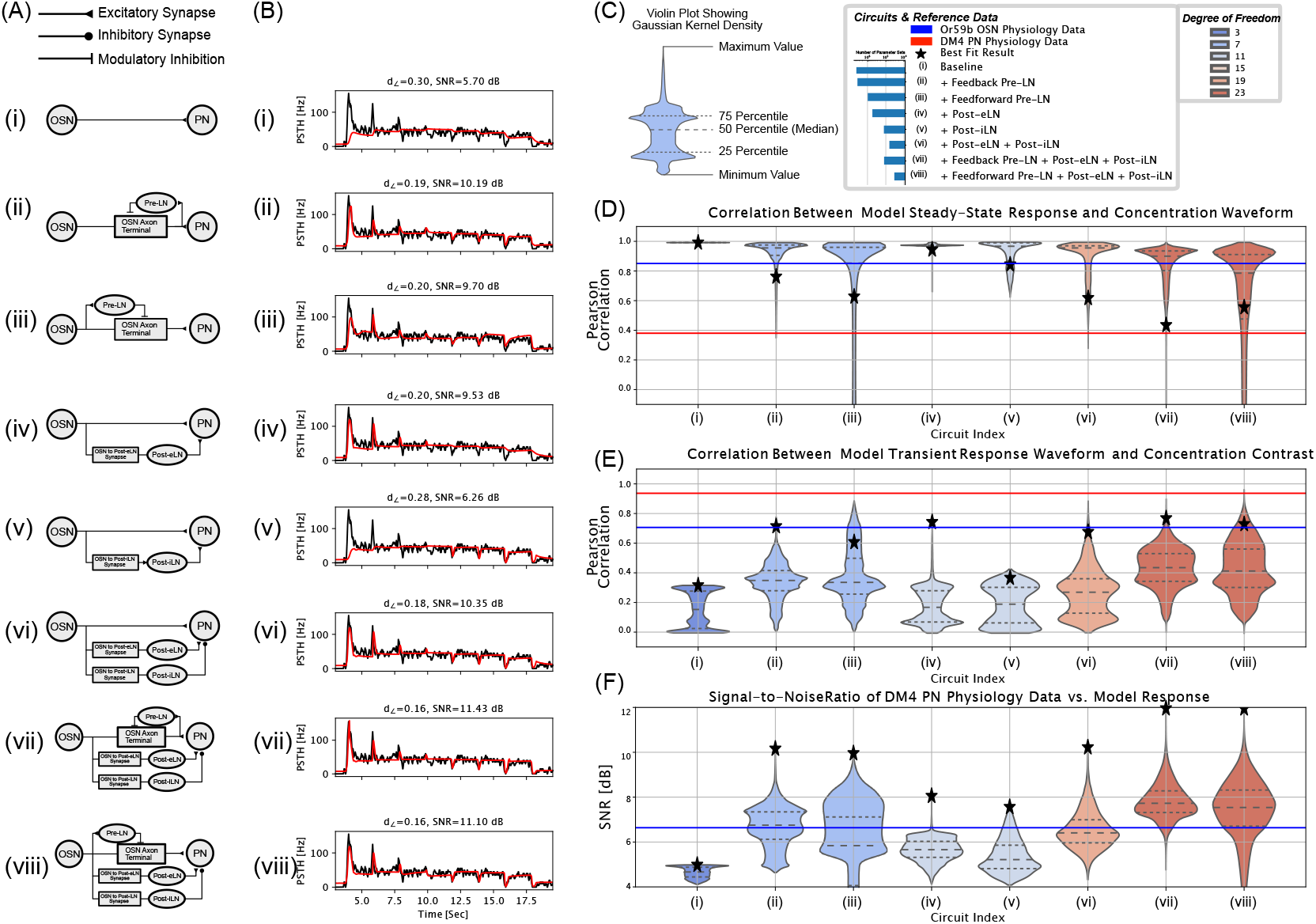
Best fit SNR and distribution comparisons between temporal odor signal processing of single channel circuit architectures (Or59b OSN and DM4 PN). (A) Circuit architectures. (B) Responses of best fit models. Other metrics associated with the best fit results are shown as black stars in (D-F). (C) Reference for violinplot, legend for model and color reference for degree of freedom (number of free parameters) of each model. (D) Sample distribution of the correlation between the steady-state responses and concentration waveform over the parameter space for each of the 8 models. The correlations between the steady-state response and the concentration waveform for DM4 PN and Or59b OSN physiology data are shown, for reference, in red and blue lines, respectively. (E) Sample distribution of correlation between the transient responses and concentration contrast over the parameter space for each of the 8 models. The correlations between the steady-state response and the concentration waveform for DM4 PN and Or59b OSN physiology data are shown, for reference, as red and blue lines, respectively. (F) Sample distribution of Signal-to-Noise Ratio between physiology recording data of DM4 PN and model responses. For reference, the Or59b OSN physiology data is linearly scaled to achieve the best SNR against the DM4 PN physiology data. The SNR between the scaled OSN response and PN response is shown in blue.

In addition to the architecture, full specification of each circuit requires model definition and parameterization of individual components. From the modeling perspective, the Calcium Feedback Loop of the Odorant Transduction Process (OTP) model of the OSN produces a transient response similar to that observed in the PN physiology data [2]. As such, the OTP model serves as a starting point for circuit component models of the Antennal Lobe.

We modeled the Pre-LN inhibition of the OSN Axon-Terminal similar to the inhibition exerted by the calcium channel of the OTP. Here the spike train of Pre-LN multiplicatively inhibits the neurotransmitter release of the OSN Axon Terminal (see Methods). The OSN-to-PostLN synapse model expands upon the calcium feedback loop to provide the flexibility to strongly respond to either input onset and offset based on choice of parameters. Finally, all other synapse models are assumed to be ionotropic and all biophysical spike generators noisy Connor-Stevens point neurons (see [2] and Methods). Together, the combined number of parameters for all components determine the degree of freedom of each circuit, an example of which is shown in **Table.**9b.

Given full circuit specifications (architecture and model parametrizations), we sought to optimize the parameters of each of the 12 circuits by minimizing the *L*_2_ distance between the temporal response of each circuit PSTH and the DM4 PN PSTH from physiology recordings. Due to the high number of parameters for the more complex circuits (e.g., 23 in the most complex circuit), we utilized a 2-step optimization procedure (see **Section** 4.3 in Methods) that combines an initial random sampling of the parameter space and a fine-tuning global optimization step. The temporal response of the circuit models with the lowest *L*_2_ distances (best fit) are shown in **Fig.**3(B) (see **Fig.**12 for results of all 12 models).

In addition to determining the best-fit circuits via the 2-step optimization procedure, we also sought a global view of the objective function landscape. Such global perspective enables comparison between the functional capabilities inherent in the circuit topology, rather than a specific parameterization. To that end, for each circuit, we randomly sampled at least 10^6^ parameters, and stimulated in excess of 5 × 10^8^ circuit/parameter combinations. Collecting results from each parameter choice, we computed the sample distribution of the SNR for all circuits, each drawn as a violin-plot in **Fig.**3(F) (see **Fig.**3(C left) for an explanation of the violin-plots). Note that that circuit name and circuit degree of freedom are detailed in the legends in **Fig.**3(C). Additionally, we computed the sample distributions of the correlation between the steady-state response and concentration waveform (**Fig.**3(D)), and the transient response and concentration contrast (**Fig.**3(E)). For reference, the same procedure is repeated for DM4 PN PSTH and Or59b OSN PSTH, and the resulting correlation values are shown in red and blue in **Fig.**3(D,E), respectively.

Referencing the results in **Fig.**3(B), we first compared the best fit results of each circuit in terms of 1) the Signal-to-Noise Ratio (SNR) between the circuit response and the PN physiology data and 2) the contrast boosting performance in terms of the response amplitudes to positive and negative concentration contrasts. Referencing the result of the comparison summarized in **Table.** 1, we note that:

- the Baseline circuit (model (i)), with no LNs, results in the lowest SNR, and fails to respond to ON/OFF odorant transients;
- the addition of Post-eLN (in model (iv)) improves the SNR significantly by 3.83 dB above that obtained with the baseline circuit and captures the odorant onset transient;
- the addition of Post-iLN (in model (v)) improves the SNR moderately by .56 dB above that obtained with the baseline circuit and captures the odorant offset transient;
- the addition of Pre-LN (in either feedback or feedforward configurations in model (ii) or (iii)) significantly improves the SNR (by 4.49 dB and 4 dB, respectively), and captures both the onset and offset odorant transients.

**Table 1:** Comparison of best fit results of each of the temporal circuits in **Fig.**3.

| Model Index | (i) | (ii) | (iii) | (iv) | (v) | (vi) | (vii) | (viii) |
| --- | --- | --- | --- | --- | --- | --- | --- | --- |
| SNR [dB] | 5.7 | 10.19 | 9.7 | 9.53 | 6.26 | 10.35 | 11.43 | 11.10 |
| Ranking by SNR | 8 | 4 | 5 | 6 | 7 | 3 | 1 | 2 |
| Odorant Transient Captured | N.A. | ON & OFF | ON & OFF | ON | OFF | ON & OFF | ON & OFF | ON & OFF |

Next, we compared the sample distributions shown in **Fig.**3(D-F), particularly the median as well as the 25/75 quantiles indicated as horizontal dashed lines within each violin-plot. Note that, for each circuit architecture, the correlations of the best fit results are marked with black stars in **Fig.**3(D-F). From **Fig.**3(F), we observe that the median SNR across parameterizations of each circuit follows the ranking of the best fit results shown in **Table.** 1. We observe that, models (ii-iii) achieve significantly higher SNR across parameterization than models (iv-v), and are comparable to the SNR achieved by model (vi). This indicates that the Pre-LN is critical in capturing the PN overall temporal response, while the simultaneous introduction of Post-eLN and Post-iLN is necessary to significantly improve the circuit’s odor signal processing capability. On the other hand, the combination of both pre-synaptic and post-synaptic LNs resulted in the highest overall SNR. These observations are supported in **Fig.**3(D), where the level of concentration-invariance is comparable between circuit (ii, iii) and circuit (vi), while a significant portion of parametrizations of circuit (vii, viii) lead to a concentration dependency level similar to that observed in the PN data. Similarly, in **Fig.**3(E), circuits (ii,iii) consistently out-perform (iv,v) in capturing the concentration contrast during transient response, and are comparable to the combined Post-eLN+Post-iLN circuit in (vi). As before, the model with the highest degree of complexity, circuits (vii, viii) show the best contrast boosting capabilities according to both the highest median as well as 75 percent quantile values, with the maximum correlation of model (viii) reaching the level observed in the PN data.

In conclusion, we have demonstrated that for the single channel temporal response of the Antennal Lobe circuit, the addition of a presynaptic local neuron (Pre-LN) significantly improves both concentration invariance and contrast boosting for both feedback and feedforward circuit configurations. Moreover, the introduction of both Post-eLN and Post-iLN is required for contrast boosting. The Post-LNs can further boost the contrast at the output of the Pre-LNs circuits without sacrificing on concentration-invariance.

### 2.2 Spatio-Temporal Presynaptic Inhibition Recovers Odorant Identity

While the single-channel circuits discussed in the previous section showed that Pre-LNs and Post-LNs support *concentration invariance* and *contrast boosting*, they only explain *temporal* features of the single channel AL odor signal processing. As odorant identities are defined *spatially* (via binding and dissociation rates across receptor types), full characterization of the AL requires analyzing multi-channel circuits involving all 23 channels with known affinity rates [25, 2].

As before, we exhaustively explored and compared the *spatio-temporal* architectures of the AL circuit that are differentiated by their innervation patterns on both the presynaptic and postsynaptic sides of the OSN-to-PN synapse. On the presynaptic side, we considered 5 circuit architectures with: 1) no Pre-LNs **Fig.**4(A[i]), 2) Pre-LNs innervating individual glomeruli in feedback or feedforward configurations **Fig.**4(A[ii,iii]), and 3) Pre-LNs innervating all glomeruli in feedback or feedforward configurations **Fig.**4(A[vi,vii]). On the postsynaptic side, the addition of Post-eLNs and Post-iLNs to circuits with local Pre-LN inhibition leads to the models in **Fig.**4(A[iv,v]), and likewise for circuits with global Pre-LN to the models in **Fig.**4(A[viii, ix]). Three additional configurations of Post-LNs were also explored (no Post-LNs, only Post-eLNs, only Post-iLNs). Hence, combining all configurations of presynaptic and postsynaptic connectivites, we compared a total of 20 different spatio-temporal circuit architectures, 9 of which are shown in **Fig.**4(A) (see **Fig.**14 for a comparison between all 20 models.)

**Figure 4:**
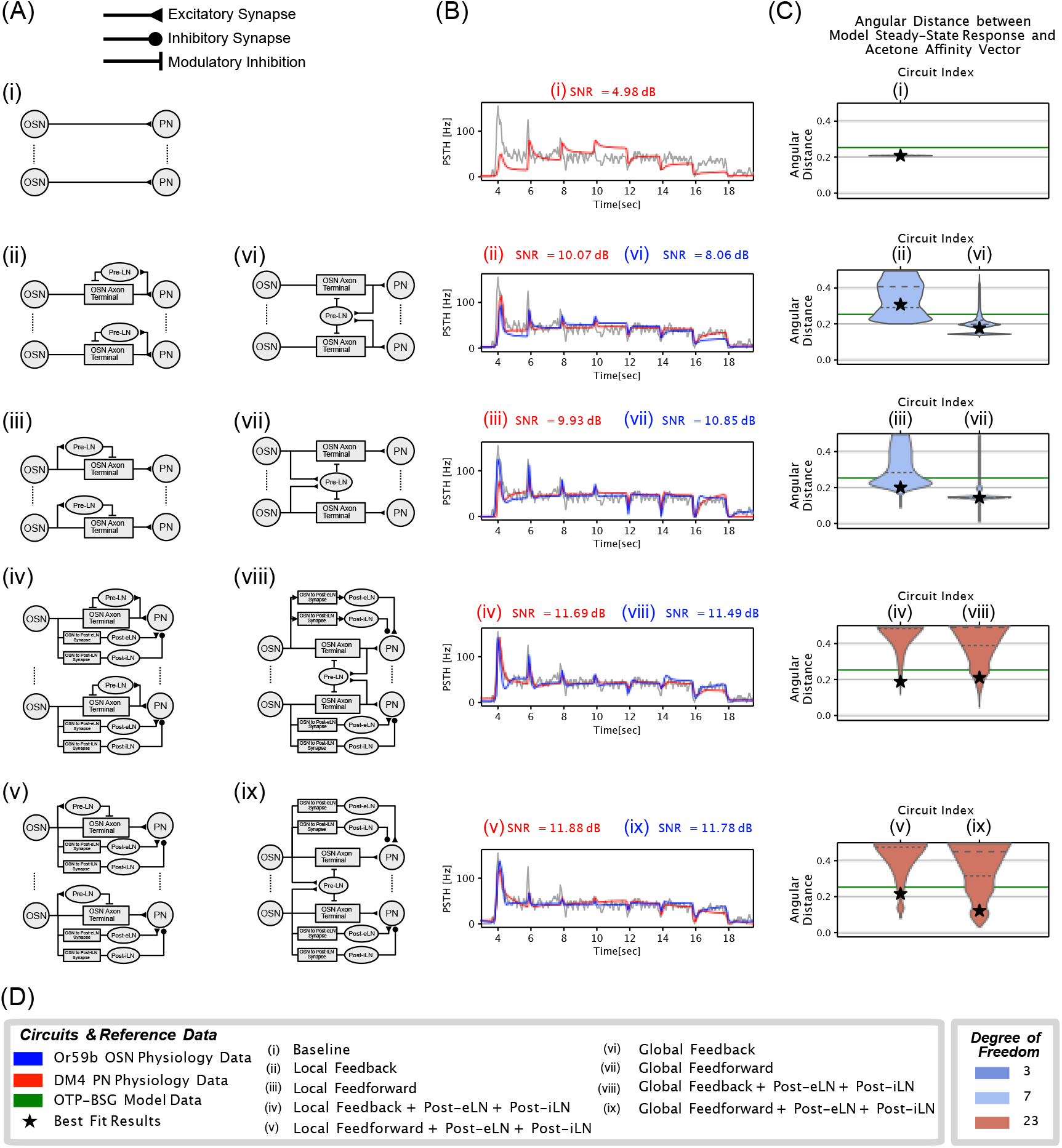
Global Feedforward/Feedback models of the Antennal Lobe outperform Local Feedforward/Feedback models across parameterization. (A) Circuit architectures, refer to (D) for names associated with each architecture. (B) DM4 PN PSTH of best fit results of each circuit. Other metrics associated with the best fit results are shown as black stars in (C). (C) Average distance between spatio-temporal model responses and Acetone odorant affinity vector. The distance between steady-state response of the OTP-BSG model and Acetone odorant affinity vector is shown in green for reference. (D) Legend for circuit architectures and degree of freedom.

While each channel in the spatio-temporal circuits receive odorant input with different binding and dissociation rates, models describing the same circuit components across channels were restricted to share the same parameter values. As such, the total degrees of freedom for the spatio-temporal circuits **Fig.**4(A[vi-ix]) is the same as that of their temporal counterparts **Fig.**4(A[ii-v]). For circuits with global Pre-LN inhibition, e.g., Global-Feedforward in **Fig.**4(A[vii]), all OSN-to-PreLN synapses share the same parameter values, and the dendritic process of the Pre-LNs was modeled as the linear sum of synaptic currents across all synapses. As no additional parameters were introduced, the Global-Feedforward circuit has the same degree of freedom as the Local-Feedforward circuit.

From an optimization standpoint, we sought to compare AL circuits both temporally and spatially. In particular, the spatio-temporal objective function (see Methods) combines the *L*_2_ distance between DM4 PN PSTHs of circuit models and physiology recordings, and the angular distance between PN steady-state response vectors and the odorant affinity vector [**b**]*_ron_*/[**d**]*_ron_*. The latter spatial objective is motivated by the hypothesis that the concentration-invariance of the DM4 PN response extends across all AL channels. As the odorant identity is represented by the affinity vector in steady-state, the objective is the angular distance between the PN steady-state response and the odorant affinity vector. The latter is minimized when the two vectors are, up to a scaling factor, identical.

Finally, the spatio-temporal optimization problem minimizes the sum of the two objectives with a weighting factor *γ* > 0 (see Methods for details), where *γ* = 0 results in the same objective for the temporal circuits (see **Fig.** 15 for the influence of *γ* on the DM4 PN PSTH of AL circuit models.) In **Fig.**4, we chose *γ* = 5 and applied the same 2-step optmization procedure and random sampling of parameter spaces as before.

From the standpoint of temporal dynamics, as the parallel processing circuit architectures in **Fig.**4 (A[i,ii,iii,iv,v]) are indistinguishable from their single-channel counterparts along the DM4 channel in (**Fig.**3 (A[i,ii,ii,vii,viii])), we expected identical (up to random parameter sampling) circuit response characteristics. Indeed, for both the best fit results and sample distribution across parametrizations, parallel processing circuits exhibit similar SNRs (**Fig.**4(B[i,ii,iii,iv,v]) and **Fig.**14(E[i,v,ix,iv,viii,xii])) with their temporal counterparts in **Fig.**3(B[i,ii,ii,vii,viii]) and **Fig.**3(F[i,ii,ii,vii,viii]). The similarities are also observed when measuring the degree of concentration-invariance and contrast-boosting along the DM4 channel as shown in **Fig.**14(C,D).

From the standpoint of spatio-temporal dynamics, comparing the angular distances between PN steady-state responses and the affinity vectors for the parallel processing circuits, we note that the Local-FF (**Fig.**4(C[ii])) and Local-FB (**Fig.**4(C[iii])) circuits exhibit some, albeit limited improvement over the baseline (**Fig.**4(C[i])) with the lowest angular distance achieved by the Local-FF model at 0.18 as compared to 0.2 for the baseline circuits. The addition of Post-LNs in **Fig.**4(C[iv,v]) resulted in a similar limited improvement in odorant recovery performance.

However, as shown in **Fig.**4(C[vi, vii]), circuits with global inhibition from Pre-LN (Global Feedback, Global Feedforward) show drastic improvement of odorant recovery as compared to their circuits with local Pre-LN inhibition, without a significant decrease in the overall SNR along the DM4 channel (see **Fig.**4(E[vi, vii])). Notably, the contrast boosting capability of the Global Feedback model (**Fig.**14(D[xiii])) is greatly reduced from its local counterpart (**Fig.**14(D[v])), which is improved by the addition of Post-LNs (**Fig.**14(D[xvi])).

In conclusion, by extending the temporal AL circuits to their spatio-temporal counterparts, and introducing circuits with global Pre-LN inhibition, we have shown that odorant identity recovery requires global Pre-LN inhibition. Additionally, we have shown that AL circuits with global Pre-LN inhibition exhibit similar temporal responses along the DM4 channel, suggesting that the improvement in odorant recovery obtained by changing local inhibition to global inhibition does not lead to a significant decrease in temporal processing performance of the circuits.

### 2.3 Modeling the Antennal Lobe as a Spatio-Temporal Divisive Normalization Processor

As shown in previous sections, Local Neurons in the Antennal Lobe form the neural basis for capturing important features of odor signal processing. To understand the functional logic of the Antennal Lobe, we sought an algorithmic description of the presynaptic and postsynaptic processing in the AL circuit that is enabled by the Pre-LNs and Post-LNs.

Comparison between the models along the Pre-LN, Post-eLN and Post-iLN pathways (see also **Fig.**9a(a) (left) and **Fig.** 16a (left)) revealed a common feature. Given input signal *v*(*t*), models of all LN pathways can incorporate the following equations:

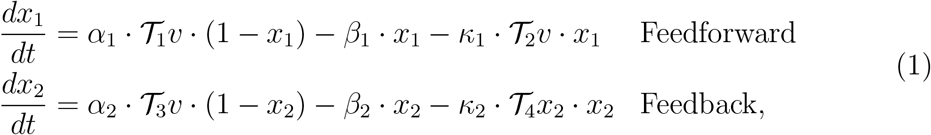

where *x*_1_(*t*) corresponds to the normalized neurotransmitter concentration in the OSN Axon-Terminal given feedforward inhibition of Pre-LN, and the normalized conductance in the OSN-to-PostLN synapse. *x*_2_(*t*) corresponds to normalized neurotransmitter concentration in the OSN Axon-Terminal given feedback Pre-LN inhibition. Additionally, 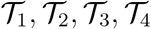 denote processors of the input signal *u*(*t*) (or state variable *x*_2_(*t*) in the feedback case). For an OSN Axon-Terminal with Feedforward Pre-LN inhibition, for example, 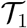 is the identity operator, and 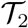 captures the non-linear transformation mapping the OSN spike train into the Pre-LN spike train (see Methods for full model descriptions). We note that the critical point (steady-state) solution for **Eq.**(1) is given as (assuming that the limit exists)

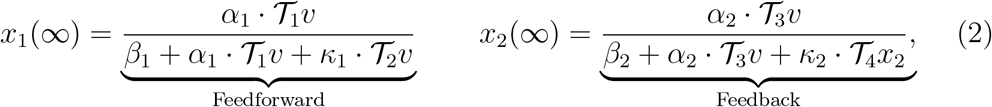

which functionally reminds us of the Divisive Normalization Processor proposed in [24]. Since the models of the Antennal Lobe operate on the circuit level described as a dynamical system, we refer to the model in **Eq.**(1) as the *differential* DNP, and the model in [24] as the *convolutional* DNP (see **Fig.**5(A2 top)).

**Figure 5:**
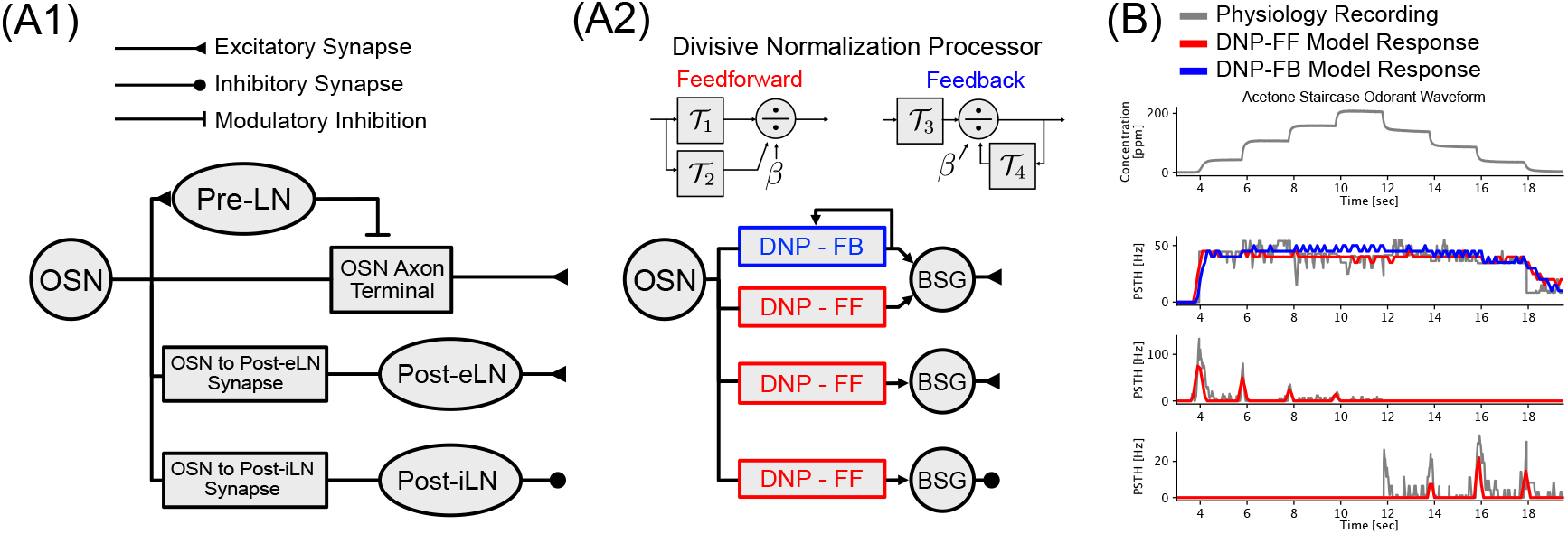
Temporal Divisive Normalization enables concentration invariance and contrast boosting for both onset and offset concentration transients. (A1) Parallel processing circuit architecture of Local-FF + Post-eLN + Post-iLN. (A2) Each branch in (A1) can be replaced by an equivalent Divisive Normalization Processor (DNP). The feedforward Pre-LN pathway in (A1) can be replaced by Feedforward (red) DNP, while the feedback Pre-LN pathway (not shown) can be replaced by Feedback (blue) DNP. (B) Input and output relationships for each of the three branches shown in (A). Odorant concentration waveform and DM4 PN physiology responses are shown in grey, DNP-FF model responses are shown in red, DNP-FB model response for Pre-LN pathway is shown in blue. (B top to bottom) Acetone staircase concentration waveform; steady-state PN response showing *concentration invariance*; transient PN response showing *contrast boosting* of ON concentration contrast; transient PN response showing *contrast boosting* of OFF concentration contrast.

Since the Local Neuron pathways (Pre-LN inhibition of OSN Axon-Terminal, OSN-to-Post-eLN synapse, OSN-to-Post-iLN synapses) can be modeled as differential DNPs, the temporal AL circuit shown in **Fig.**3(A[viii]) reduces to 3 parallel sub-circuits (**Fig.**5(A1)) each modeled by a differential DNP (**Fig.**5(A2)) in cascade with a Biophysical Spike Generator (modeled here as Connor-Stevens point neuron). Although the Pre-LN pathway in **Fig.**5(A1) does not incorporate a BSG, we added for symmetry reasons a BSG to DNP-FF[I] in **Fig.**5(A2). A removal of the BSG in DNP-FF[I] does not result, however, in a qualitative change in the encoding characteristics along this pathway (data not shown). As the comparative analysis of AL circuits in **Fig.**3 suggest that the Pre-LNs and the Post-LNs contribute differently to the PN response characteristics, we hypothesize that the three DNPs function independently to capture concentration invariance, ON and OFF contrast boosting, respectively (**Fig.**5(B)).

To verify this hypothesis and show that DNPs can indeed capture these features of PN responses, we first decomposed the DM4 PN response into concentration invariant steady-state and contrast boosting ON and OFF transient responses as shown in **Fig.**5(B grey) (see Methods for details). We then optimized the three Feedforward DNPs in cascade with three Connor-Stevens BSGs (DNP-FF shown in red in **Fig.**5(A2)) each with the objective function given as the angular distance between model BSG PSTH (**Fig.**5(B red)) and physiology PSTH. Applying the two-step optimization procedure, we conclude that the best fit results for DNP-FFs can indeed separately capture the three response features. Additionally, in agreement with findings in **Section** 2.1, the Feedback DNP is also able to capture concentration invariance along the Pre-LN pathway as shown in **Fig.**5(B blue).

Similar to the temporal DNP in **Eq.**(1), spatio-temporal models of the OSN Axonn-Terminal can be described by *Global Differential DNP* equations:

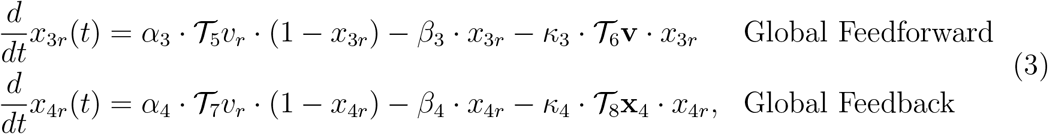

where *r* = {1,…, *R*} is the index for channels in the spatio-temporal AL circuit, *v_r_*(*t*) represents input in the *r*-th channel, 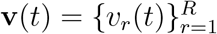 represents multi-dimensional input across channels, *x*_3*r*_(*t*)(*x*_4*r*_(*t*)) represents output signal of Feedforward (Feedback) DNP in *r*-th channel, and 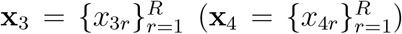 represents multi-dimensional output signals of Feedforward (Feedback) DNP across channels.

As in **Eq.**(2), we note that the critical point solutions of the Global Differential DNP are given as:

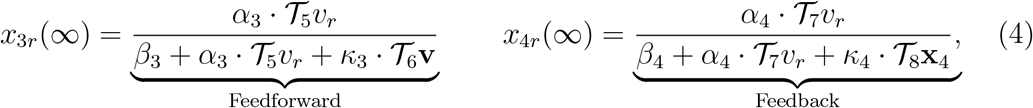

Given that the odorant transduction process involves a multiplicative coupling between the affinity vector and the odorant concentration

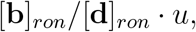

the spatio-temporal divisive normalization provides, in steady state, a normalized odorant affinity vector without prior specification of the odorant identity or concentration profile [11]. Furthermore, as shown in **Fig.**4(A), we note that the differential DNPs discussed above can also be extended to their spatio-temporal counterparts (Global DNP-FF and Global DNP-FB in **Fig.**6(A2), see Methods for model details). As shown in **Fig.**6(B[i,ii]), both the Global DNP-FF and Global DNP-FB circuits are able to capture concentartion invariance along the DM4 channel. Furthermore, the spatio-temporal responses in **Fig.**6(B[iv,v]) recovers the Acetone affinity vector with an average angular distance between circuit response and affinity vector of 0.13 for Global DNP-FB and 0.12 for Global DNP-FF, respectively. For reference, the average angular distance between the OSN response and the affinity vector is 0.467.

**Figure 6:**
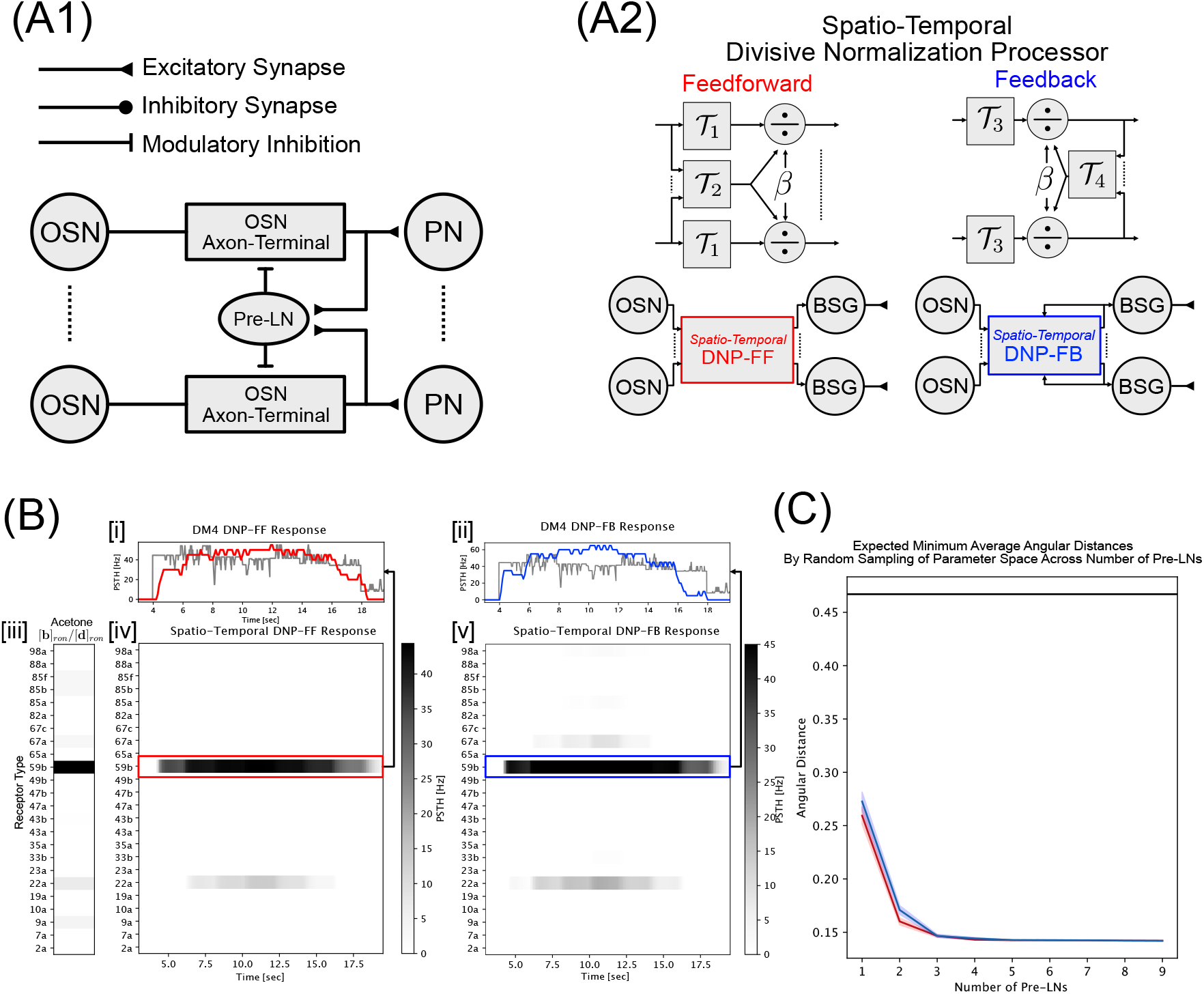
Spatio-temporal Divisive Normalization along the Pre-LN pathway recovers odorant identity. (A1) Circuit architecture of Global-FF as in **Fig**.4(A[vi]). (A2) The feedback Pre-LN inhibition in (A1) can be replaced by an equivalent Global Feedforward (red) DNP while the feedback Pre-LN inhibition (not shown) can be replaced by a Global Feedback (blue) DNP. (B) Spatio-temporal input and output relationships of Global DNP-FF and Global DNP-FB. [i] Global DNP-FF temporal response along DM4 channel correspoding to OR59b receptor type. [ii] DM4 Global DNP-FB temporal response. [iii] Acetone affinity vector representing odorant identity. [iv] Spatio-temporal Global DNP-FF response for all 24 channels with known affinity rates. [v] Spatiotemporal Global DNP-FB response for all 24 channels with known affinity rates. (C) Expected minimum average angular distance between model PSTH (Global DNP-FF in red, Global DNP-FB in blue) and Acetone affinity vector. The standard error in the minimum average angular distance is shown as shaded red/blue regions around each solid line. For reference, average angular distance between OSN response and affinity rate is shown as the horizontal black line around *d*_∠_ = 0.467.

To show that the DNPs can robustly recover odorant identity across parameterizations, we randomly sampled 10^6^ parameter sets for both the Global DNP-FF and the Global DNP-FB, and found that over 82% and over 85% of the randomly sampled parameter sets for Global DNP-FB and DNP-FF show a reduction in average angular distance from the OSN responses. These results support the hypothesis that the global Pre-LN inhibition robustly recovers odorant identity.

Given that each neuron type in the *Drosophila* connectome is instantiated by multiple individual neurons [21], we further asked the question of what is the expected odorant recovery performance if the Pre-LN pathway is comprised of *N* DNPs parameterized at random? Following a method similar to boostrap sampling [26] we randomly sampled *N* (*N* = {1,2,…, 10}) different parameter sets of the Global DNP-FF and Global DNP-FB and recorded the *minimum* average angular distance amongst the samples. Repeating this processing 1000 times for each value of *N*, we estimate the mean and standard error of the minimum average angular distance achieved by *N* DNPs describing the Pre-LN pathway. As shown in **Fig.**6(C), we observe that, even with a single Pre-LN, the expected angular distance around 0.27 for both Global DNP-FF and DNP-FB is significantly lower than that of the OSN. Additionally, at 3 Pre-LNs and beyond, the angular distance converges to below 0.15, a value that is consistent with the average angular distance achieved by the best performing Global-FF and Global-FB AL circuit models shown in **Fig.**4(F[vi,vii]).

In conclusion, we show that the AL circuit can be described as three parallel DNPs modeling Pre-LN, Post-eLN and Post-iLN pathways, which are indepedently tuned capture concentration invariance and ON/OFF contrast boosting PN response features, respectively. More importantly, the separation into 3 DNPs prompted us to hypothesize that the AL implements an *ON-OFF odorant identity recovery* processing paradigm. Under such paradigm, the Post-eLNs and Post-iLNs signal changes its receptor responses across channels, while the Pre-LNs extract the odorant identity vector for downstream odorant recognition and associative learning.

### 2.4 The Antennal Lobe is a Robust ON-OFF Odorant Identity Recovery Processor

The theoretical analysis in the previous section prompted us to hypothesize that the functional logic of the AL circuit is *ON-OFF odorant identity recovery*. While the lack of spatio-temporal experimental data limits the biological validation of such a hypothesis using AL PN recordings, a massive computational evaluation of the robustness of ON-OFF detection and odorant identity recovery is a tractable alternative.

To robustly signal ON/OFF event timing, the Post-LN pathways have to respond reliably for different odorant concentration waveforms and amplitudes. Using 3 odor-ant waveform profiles with different concentration contrasts (step, ramp, parabola, **Fig.**7(A1,B1,C1)) [23], we evaluated the corresponding DNP-FF responses along the Post-eLN and Post-iLN channels for concentration amplitude values ranging from 1 to 10,000 ppm (**Fig.**7(A2,B2,C2)). To quantify the input dependency of the responses, we measured the amplitude and timing of the peak responses of both Post-eLN and Post-iLN pathways **Fig.**7(A3,B3,C3). As depicted in **Fig.**7(A3, left), the peak Post-eLN (shown in red) and Post-iLN (shown in blue) spike rates remain relatively constant for odorant concentration amplitude values above 2,000 ppm, while the OSN spike rate (shown in black) increased by almost 50% from ~ 150Hz at 2,000 ppm to around ~ 220 Hz at 10,000 ppm. These results suggest that the responses of Post-eLNs and Post-iLNs are robust to wide changes of concentration amplitude values. Additionally, the timing of the peak responses is remarkably consistent across concentration levels as shown in **Fig.**7(A3, right), indicating that the Post-LNs provide ON/OFF timing information regardless of odorant concentration levels. Furthermore, while the Post-LN responses to ramp and parabola waveforms are weaker due to the significantly lower contrast levels, both the onsets and offsets still trigger strong Post-LN responses at higher odorant concentration levels (**Fig.**7(B3, C3)). Thus, the ON-OFF event detector is robust to both concentration amplitude levels and waveform profiles.

**Figure 7:**
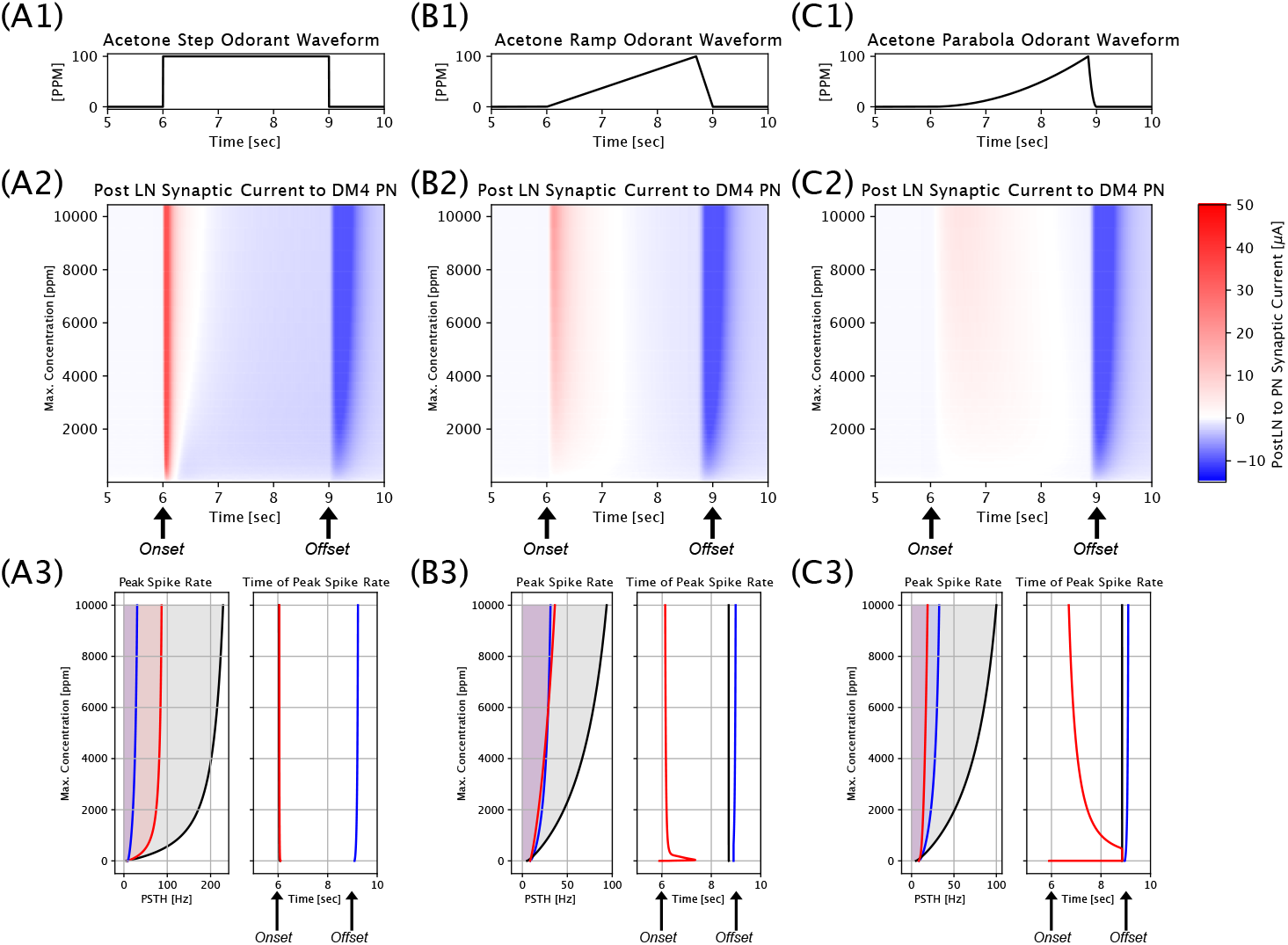
Postsynaptic Local Neurons (PostLNs) robustly encode odorant on/off-set timing information. (A1) Step Acetone waveform input to OSNs. (A2) Synaptic current from Post-eLN and Post-iLN to DM4 PN across concentration levels from 1 to 10^4^ ppm for step concentration waveform. Post-eLN-to-PN synaptic current is shown in red and Post-iLN-to-PN synaptic current is shown in blue. (A3) Amplitude and timing of the peak Post-eLN (red) and Post-iLN (blue) responses in comparison to OSN (black). (B1-B3) Same as (A1-A3) for ramp concentration waveform with smaller onset/offset concentration contrast. (C1-C3) Same as (A1-A3) for parabola concentration waveform with the smallest onset/offset concentration contrast.

Upon event-detection in the Post-LN pathways, the AL recovers the odorant identity across concentration levels. To evaluate the robustness of odorant identity recovery, we compared the steady-state PN responses of the spatio-temporal circuits with the affinity vectors of each of the 110 odorants with known affinity vectors [2, 25]. For each of the odorants we used step concentration waveforms with an amplitude ranging from 10 ppm to 1,000 ppm. The resulting distance measures are shown in **Fig.**8(B) as heatmaps. Here each row corresponds to a specific concentration level (in log scale), and each column corresponds to an odorant identity. Note that, across circuits, for each odorant (along each column), the angular distance is highest (indicating worst recovery) at very low/high concentration levels, and is lowest at intermediate concentration levels. The histograms of angular distances shown as heatmaps in **Fig.**8(B) are displayed as violin plots in **Fig.**8(D). The results depicted clearly show that the Global Feedback and Global Feedforward models achieve significantly better odorant recovery than the other circuit models considered. Finally, to isolate the computation performed by the Antennal Lobe from that by the Antenna, we plotted the angular distance between (PN steady-state response, affinity vectors) versus (OSN steady-state response, affinity vectors) in **Fig.**8(C). Since the degree of odorant recovery is inversely proportional to the angular distance, an *improvement* from Antenna response would by reflected in **Fig.**8(C) as data below the diagonal line. Importantly, as shown in **Fig.**8(C[iv, v]), the global inhibition models show an uniform improvement of odorant identity recovery, with minimum angular distances reaching 0, indicating perfect odorant identity recovery. Circuits with local Pre-LN inhibition, on the other hand, do not show such uniform improvement of odorant recovery (**Fig.**8(C[i-iii]).

**Figure 8:**
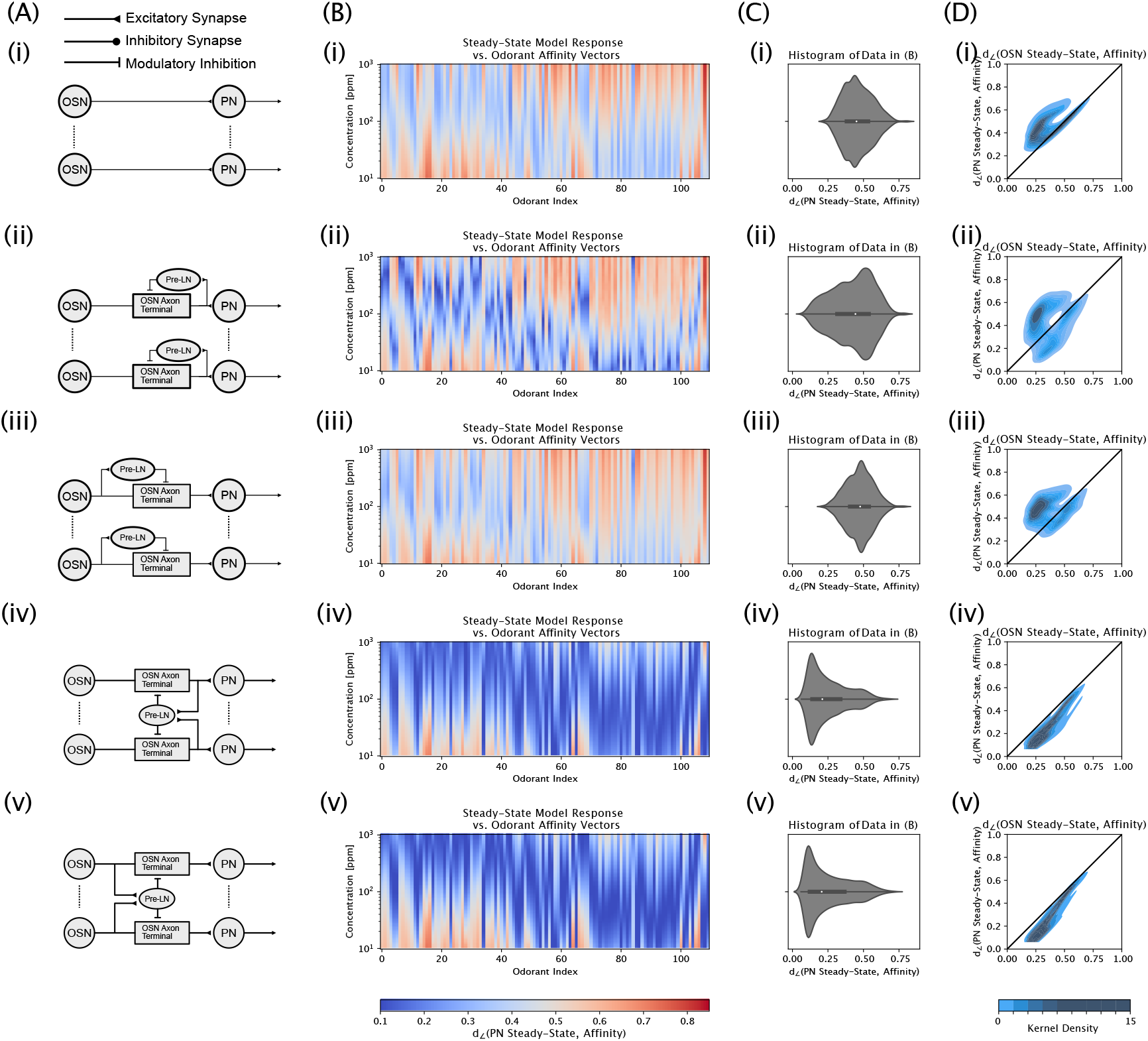
Recovery of mono-molecular odorant identities by spatio-temporal response of the Antennal Lobe for 110 odorants with known estimated affinities across concentration levels. 5 models of the spatio-temporal processing of tspahe Antennal Lobe is considered for the odorant identity reconstruction evaluation, each shown as a row indexed [i] to [v]. For each model (row), the figures (column) indicate: (A) Illustration of models; (B) Angular distance between model steady-state responses and affinity vectors for 110 odorants at 10 ~ 1000 [ppm] concentration levels. (C) Histogram of data in (B) presented as violin-plots. (D) Comparison of data in (B) vs. angular distance between steady-state response of OSNs and affinity vectors (data not shown) presented as heatmap estimated using 2D kernel density estimation with gaussian kernel. Note that data below the diagonal line indicates a reduction of angular distance between model response and affinity vector over the OSN response, vice versa.

In conclusion, we computationally demonstrated that the Pre-LNs, Post-eLNs and Post-iLNs, undelying the neural basis of odor signal processing in the AL, facilitate robust PN spatio-temporal transient and steady-state responses across odorant identity and concentration waveforms. As such, they implement the functional logic of *robust ON-OFF odorant identity recovery* in the AL.

## 3 Discussion

In complex olfactory environments, *Drosophila* need to rapidly and robustly identify and react to changing odorant identities. As the encoding of odorants is characterized by the multiplicative coupling between the *odorant object identity* (semantic information) and the *concentration waveform* (syntactic information), [2], the two sources of information are confounded at the level of the Antenna [7, 8]. Single channel PN physiology data [23] show that PN temporal responses exhibit *concentration-invariance* and *contrast-boosting* properties, indicating a decoupling at the output of the AL of the odorant object identity from concentration waveform in steady-state while responding strongly to odorant concentration onset and offset in transient states [27, 28]. In the current work, we sought to study the functional logic of the Antennal Lobe that supports the observed temporal response features, and the recovery of odorant object identity from the confounding representations.

The connectome of adult [21] and larva [22] *Drosophila* AL has revealed extensive innervation by a wide variety of Local Neurons (LNs) characterized by their connectivity patterns and neurotransmitter profiles. Focusing on three types of LNs with presynaptic inhibition and postsynaptic inhibition/excitation of the OSN-to-PN synapse, we simulated all (32) circuit architectures of the AL via massively parallel program execution on Graphical Processing Units (GPUs) [29, 30]. Furthermore, for each circuit configuration, we computed empirical distributions of identity recovery across randomly sampled parameters, providing insights into the computational capabilities of model structures, rather than specific parameterizations of said models.

We found that, for temporal processing of odors along a single channel, inhibition from Pre-LNs supports *concentration-invariance* in PN responses, while Post-eLNs and Post-iLNs enhance the *contrast-boosting* across all parameterizations.

For spatio-temporal processing of mono-molecular odorants, we hypothesized that the population steady-state PN response is a concentration-invariant recovery of the odorant identity in terms of its affinity vector. We note that by recovering the odorant affinity vector, AL responses also disambiguate between different odorant stimuli, consistent with observations made by previous studies investigating a small subset of glomeruli [31, 32, 33, 34]. We found that inter-glomeruli communication with Pre-LN connecting globally to all glomeruli (in either feedback (Global-FB) or feedforward (Global-FF) configurations) is necessary to recover the odorant affinity vector at the level of the PNs.

We showed that the Antennal Lobe circuits can be algorithmically described by (differential) spatio-temporal Divisive Normalization Processors (DNPs)[24]. Specifically, the AL circuit is equivalent to 3 separate DNPs, each modeling the Pre-LN/Post-eLN/Post-iLN pathways, and extracting the concentration invariance and ON/OFF contrast boosting features in parallel and independently of each other pathway. Furthermore, consistent with previous experimental/computational results [31, 35, 32, 34, 11], the spatio-temporal DNP along the Pre-LN pathway naturally reduce the concentration dependency of the steady-state population PN response recovering the odorant object identity (semantic information) as odorant affinity vectors. As such, the separation of LN pathways suggests a feasible processing paradigm where the odorant object identity and the odorant concentration waveform are separately extracted and processed by the LN pathways.

From a robustness perspective, we showed that the Post-eLNs and Post-iLNs can act as robust event detectors by capturing odorant onset and offset timing information across concentration waveform profiles. Additionally, the Pre-LN pathway’s odorant recovery is robust across odorant object identity and concentration waveform profiles. The robustness of the LN pathway responses, and the separation of odorant object identity and concentration waveform profile along these pathways strongly suggest that the functional logic of the AL is <u>ON-OFF odorant object identity recovery</u>, enabling rapid and robust odorant object recognition and associative memory in downstream neuropils.

As the odorant object identity is characterized by the PN spatio-temporal population response, experimental verification of our models require, ideally for all glomeruli, simultaneous PN recordings. For example, while the temporal response of DM4 PN was verified against physiology data, the evaluation of the spatio-temporal AL response can only be compared against odorant object affinity vectors, and not physiology recordings. Biological verification of the spatio-temporal model, therefore, calls for massive recordings and a systematic characterization of populations of PN responses.

Additionally, while the AL circuit models compared are exhaustive for the LN cell types considered, they do not capture the full complexity of the Antennal Lobe. For example, the AL processing paradigm considered here assumes that the population response of the uni-glomerular PNs recover the odorant object identity, without a need to account for the multi-glomerular PNs. Furthermore, the LN cell-types are assumed to be either pan-glomerular or uni-glomerular, while many LNs in the AL fall in between these two extremes. Nevertheless, the circuit models presented herein can be readily extended to incorporate the additional neuron types.

## 4 Materials and Methods

In **Section** 4.1, we describe the methodology and modeling details of temporal and spatio-temporal AL circuits. In **Section** 4.2 we define concentration contrast and describe the procedures for decomposing the PN PSTH into steady-state and ON/OFF transient components. In **Section** 4.3 we define the optimization approach and procedures for evaluating the functional logic of the AL circuits.

### 4.1 DNP Models of the Architecture of the Antennal Lobe

As shown in **Section** 2.3, all LN pathways (Pre-LN, Post-eLN, Post-iLN) in the Antennal Lobe explored in **Fig.**3 and **Fig.**4 can be characterized by differential DNPs described by **Eq.**(1) and **Eq.**(3). In what follows we detail two examples of the temporal (**Fig.**9a) and the spatio-temporal (**Fig.** 10a) AL circuits, and their corresponding DNPs. All 12 temporal AL circuits and 20 spatio-temporal AL circuits considered in the current work can be similarly constructed from the examples given here.

**Figure 9:**
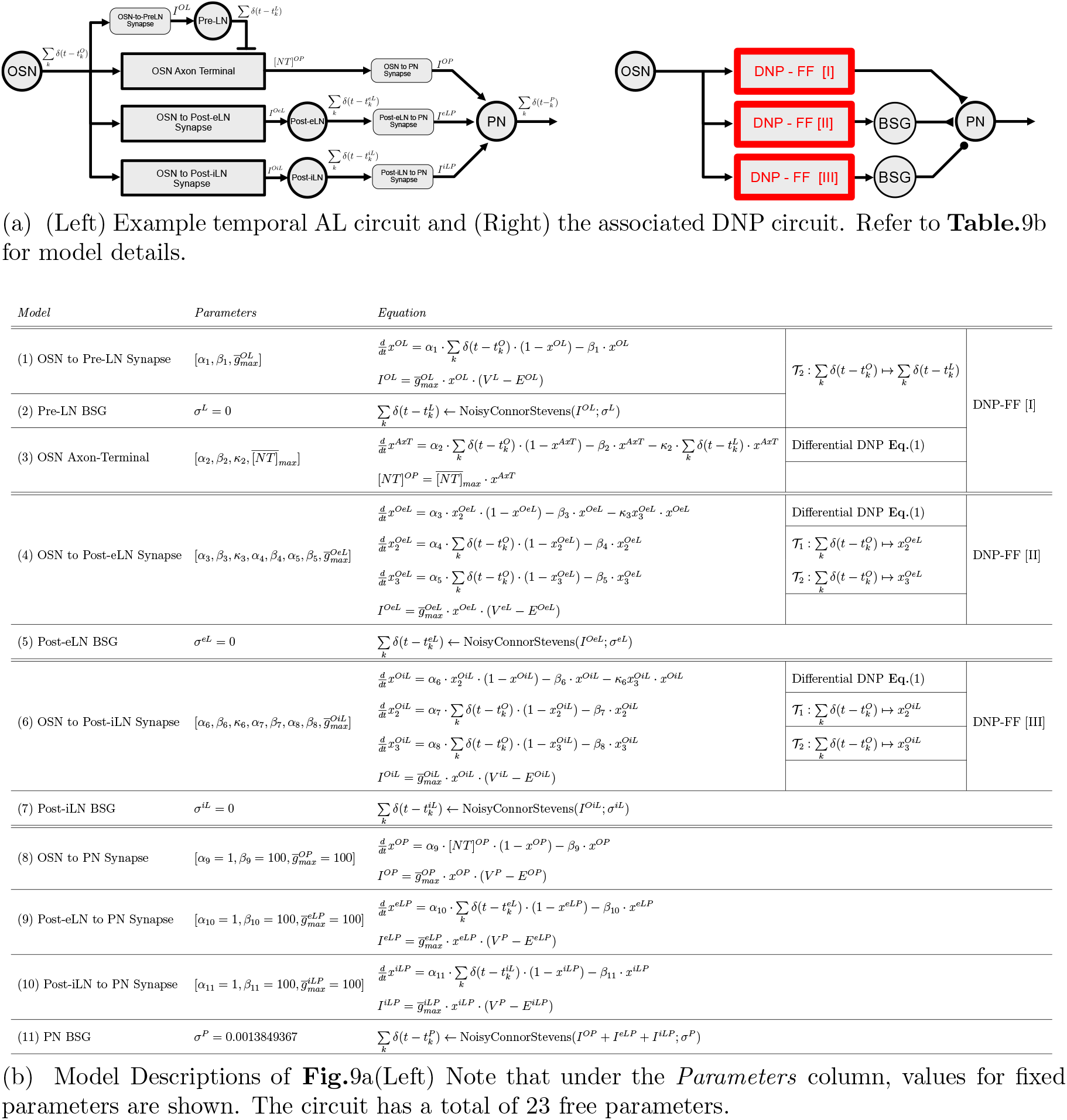
Example temporal AL circuit and associated equations.

**Figure 10:**
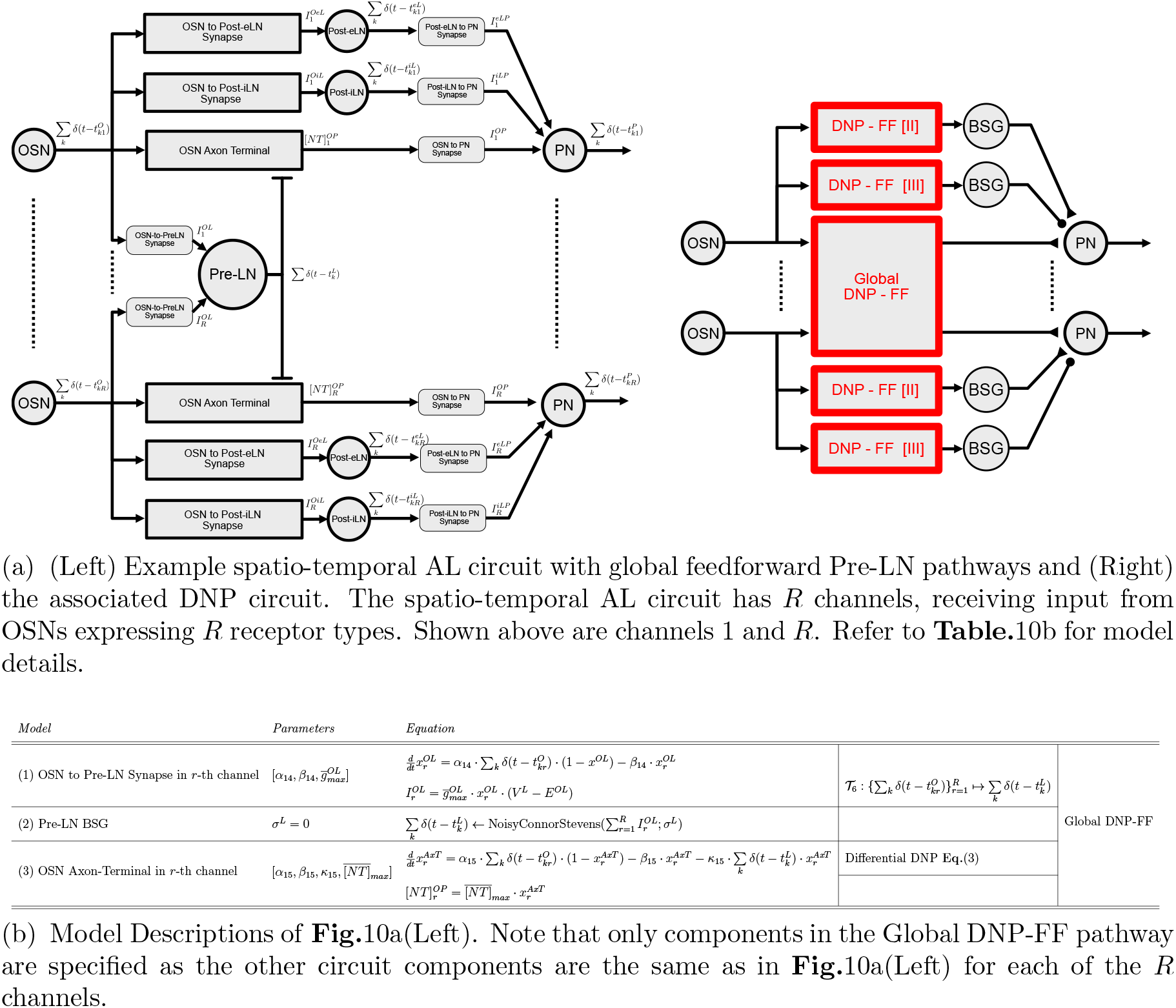
Example spatio-temporal AL circuit and associated equations.

#### Temporal Antennal Lobe Circuit

Here, we detail the models describing the temporal AL circuit with 1) local feedforward Pre-LN inhibition of the OSN Axon-Terminal, 2) Post-eLN pathway, and 3) Post-iLN pathway. Each circuit component of the example AL circuit in **Fig.**9a(left) is described by a set of differential equations detailed in **Table.**9b. Additionally, as shown in **Fig.**9a(right), the three LN pathways correspond to three feedforward temporal DNPs (DNP-FF [I-III]). For example, we note that the DNP-FF [I] in **Fig.**9a(right) describes the transformation mapping the OSN spike train 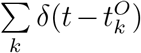 into neurotransmitter concentration [*NT*]*^OP^*, implemented by **Table.**9b(1-3). Specifically, as highlighted in **Table.**9b, the differential DNP-FF[I] (**Eq.**(1), top) is described by the OSN Axon-Terminal, while the transformation 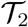 in **Eq.**(1) is described by the OSN to Pre-LN synapse and the Pre-LN BSG. The correspondence between the AL circuit and DNP-FF [II,III] are similarly highlighted in detail in **Table.**9b.

Based on the circuit in **Fig.**9a (left), a total of 6 configurations (out of 12 described in **Section** 2.1) can be generated by selectively removing one or more LN pathways following the procedures below:

1. *Pre-LN Pathway*: removing the Pre-LN pathway can be achieved by silencing the *OSN to Pre-LN Synapse*, and is implemented by removing rows (1-3) in **Table.**9b, and using 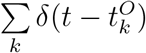 as input to the *OSN to PN synapse*.
2. *Post-eLN Pathway*: removing the Post-eLN pathway can be achieved by silencing the *OSN to Post-eLN Synapse*, and is implemented by removing rows (4,5) in **Table.**9b.
3. *Post-iLN Pathway*: removing the Post-iLN pathway can be achieved by silencing the *OSN to Post-iLN Synapse*, and is implemented by removing rows (6,7) in **Table.**9b.

To generate the 6 remaining temporal AL circuits, the Pre-LN pathway in **Fig.**9 is changed from a *feedforward* to a *feedback* configuration. Referencing the circuit architecture in **Fig.** 16a, and the corresponding model descriptions in **Table.** 16b, the *feedback* Pre-LN pathway uses the neurotransmitter concentration of the OSN Axon-Terminal [*NT*]*^OP^* as input to the *OSN to Pre-LN Synapse*.

#### Spatio-temporal Antennal Lobe Circuit

The spatio-temporal AL circuits described in **Section** 2.2 receive input from OSNs expressing a total of *R* receptor types. To simplify the notation, we denote the OSN spike train from the *r*-th channel (expressing *r*-th receptor type) as 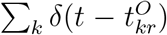, with the understanding that multiple OSNs expressing the same receptor type may innervate the same glomerulus.

The temporal AL circuit presented in the previous section can be easily generalized to their spatio-temporal counterparts. By repeating the temporal circuit in a parallel fashion for all *R* channels, 12 spatio-temporal AL circuits (out of the 20 described in **Section** 2.2) with parallel processing can be generated. As the only difference between the channels in parallel processing circuits lies at the input level (different OSN spike trains), we omit the model descriptions of these 12 circuits.

To model inter-channel communications via Local Neurons, we describe in detail a spatio-temporal AL circuit with 1) global feedforward Pre-LN inhibition of OSN Axon-Terminals, 2) local Post-eLN pathways, and 2) local Post-iLN pathways. As shown in **Fig.**10a, the three LN pathways correspond to three feedforward DNPs (Global DNP-FF, DNP-FF [I], DNP-FF [II]). However, as the Post-eLN and Post-iLN pathways are local to each channel, they are fully specified by their temporal counterparts in the previous section. As such, only the Global Pre-LN pathway is specified in **Table.** 10b.

As in the case of the Temporal AL circuits, we selectively removed one or more LN pathways in **Fig.** 10. Consequently, a total of 6 configurations (out of 20 described in **Section** 2.2) were generated from the circuit in **Fig.**9a following the procedure below:

1. *Global Pre-LN Pathway*: removing the Global Pre-LN pathway can be achived by silencing the *OSN to Pre-LN Synapses*, and is implemented by removing rows (1-3) in **Table.** 10b.
2. *Local Post-eLN Pathway*: removing local Post-eLN pathway follows the same procedure as for the temporal AL circuits, see previous section for details.
3. *Local Post-iLN Pathway*: removing local Post-eLN pathway follows the same procedure as for the temporal AL circuits, see previous section for details.

However, we note that if the Global Pre-LN pathway is removed from **Fig.** 10 (account for 2/6 configurations), the circuit reduces to a parallel processing model. As such, 4 unique configurations of the spatio-temporal AL circuits can be generated by simplifications of **Fig.** 10.

To generate the 4 remaining spatio-temporal AL circuits, the Global Pre-LN pathway in **Fig.**10 is changed from *feedforward* to *feedback* configuration. Referencing the circuit architecture in **Fig.** 17a, and the corresponding model descriptions in **Table.**17b, the *feedback* Pre-LN pathway uses the neurotransmitter concentration of the OSN Axon-Terminal 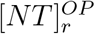 as input to the *OSN to Pre-LN Synapses*.

### 4.2 The Structure of Time Evolution of OSN and PN Responses to Odorants Waveforms

#### Concentration Contrast

We define the contrast of an odorant concentration waveform *u*(*t*) as:

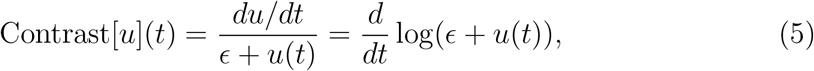

where *ϵ* is a small bias term that avoids division by zero. For concentration contrast of Acetone staircase odorant waveform as shown in **Fig.**2(A bottom), *ϵ* was set to 1.

#### Decomposing OSN and PN Responses into Steady-State and Transient Components

For odorant concentration waveforms that are piecewise constant (e.g., step, staircase), we linearly decomposed the neuron response PSTH into steady-state and transient components using the following steps (see **Fig.**2(C2) for example):

- *Extracting the odorant concentration waveform jump times*: we first computed the odorant concentration contrast using **Eq.** (5) with *ε* =1, and stored the timing when peak values are achieved as jump times.
- *Computing the steady-state component*: we then created a piecewise constant steady-state response with transition times specified by the jump times, and the amplitude specified by the average response during the 500 milisecond prior to each jump time.
- *Computing the transient component*: we created the transient response as the residual obtained by subtracting the steady-state response from the overall response.

The transient component can be further decomposed into ON and OFF sub-components, where the ON component corresponds to the non-negative transient response, while the OFF component to the non-positive response.

### 4.3 Methods of Circuit Optimization of the Antennal Lobe

#### Metrics

The concentration waveforms and output PSTHs considered in the current work are assumed to be in the space of square-integrable functions (not necessarily bandlimited). For such signals, we respectively use the *L*^2^ norm and the associated inner-product

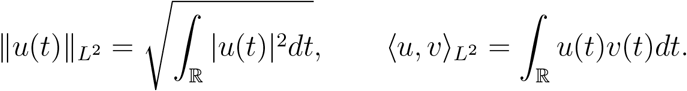

For a clean signal *u*(*t*) and the noisy signal 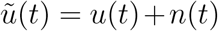, where *n*(*t*) is the noise, the SNR is given (in deciBel) as

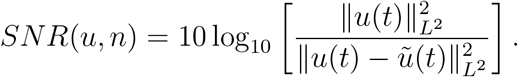

Consider two signals *u*(*t*), *v*(*t*) with temporal support [*T*_1_, *T*_2_]. The Cross-Correlation is measured as:

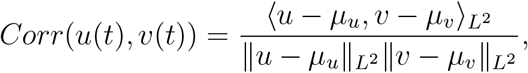

where 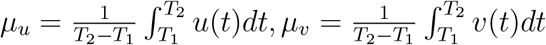 are, respectively, the average value of the two signals on [*T*_1_, *T*_2_]. To account for delay between signals *u*(*t*), *v*(*t*), Maximum Cross-Correlation can be used to robustly measure the similarity between the two signals, given as:

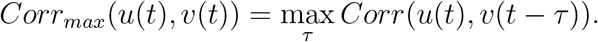

A scale-invariant pseudo-metric often used in the current work is the angular distance, defined as:

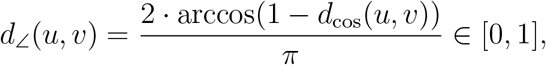

where *d*_cos_ is the cosine distance given as:

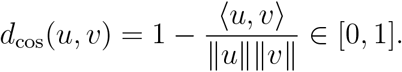

#### Optimization Objectives

For temporal models presented in **Section** 2.1, we sought to minimize the distance between PN PSTH of the models parameterized by 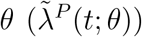 and physiology response *λ^P^* (*t*):

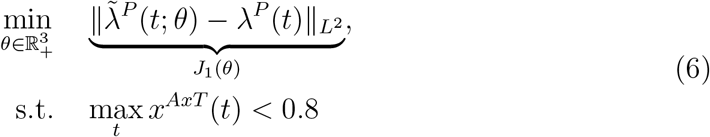

where *x^AxT^*(*t*) indicates the normalized neurotransmitter concentration of the OSN Axon-Terminal (see **Table.**9b). The constraint on normalized conductance *x^AxT^*(*t*) ∈ [0, 1] is a heuristic preventing the saturation of neurotransmitter concentration of the OSN Axon-Terminal, which ensures that concentration-invariance is achieved via the model structure, instead of an artifact of model saturation.

For spatio-temporal models of the Antennal Lobe, the overall optimization objective is described as:

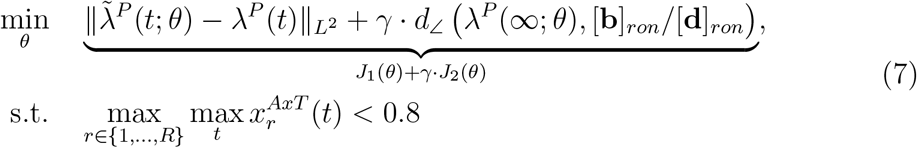

where 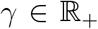 is a scaling factor, [**b**]*_ron_*/[**d**]*_ron_* is the affinity vector of the odorant identity, *λ^P^*(*t*), 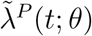 are physiology and model PN PSTH, and 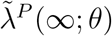 is the steady-state mode PN PSTH. For staircase concentration waveforms, 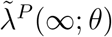 is the average PN PSTH over 500 miliseconds before jump times in odorant waveforms (see above). Similar to the temporal objective in **Eq.**(6), a constraint is placed on the maximum normalized neurotransmitter concentration of the OSN Axon-Terminal across channels and time.

In **Eq.** (7), the scaling factor *γ* controls the balancing between fitting DM4 physiology response vs. recovery of odorant affinity vector. The DM4 model response for different choices of *γ* in optimizing spatio-temporal models of the AL circuit is shown in **Fig.** 15.

#### Two-Step Optimization Procedure

Optimization of parameters for all models presented herein follow a two-step algorithm: 1) random sampling in the parameter space using Latin Hypercube Sampling [36], 2) fine tuning using Differential Evolution [37].

##### Step 1: Latin Hypercube Sampling (LHS) [36]

Latin Hypercube Sampling (LHS) is a method of generating random samples of parameter values from a multi-dimensional distribution. As opposed to random sampling (which generates samples uniformly along each dimension) or grid sampling (which generates samples at regular intervals along each dimension), LHS tries to optimize the random samples to satisfy the Latin Hypercube condition - one and only one sample is contained in each axis-aligned hyperplane. Specifically, a correlation-minimization optimization procedure was used to further randomize the initial uniform random samples in the work presented herein. The LHS routine utilized is adapted from scikit-optimize [38].

##### Step 2: Differential Evolution [37]

Differential Evolution (DE) is an evolutionary algorithm that seeks to solve a global optimization problem by iteratively improving candidate solutions within a parameter space hypercube. For the two-step optimization procedure, the constraint hypercube is formed by: 1) selecting the top 100 parameter sets found using LHS sampling, 2) computing the mean and the standard deviation of parameter values along each dimension, and 3) creating hypercube where each edge spans the mean parameter value **±**1 standard deviation. The DE algorithm then randomly recombine and replace candidate solutions sampled within the hypercube if the recombinations result in a reduction in the optimization objective. The DE optimization procedure used in the current work is an accelerated implementation of scipy [39].

## Data and Code Availability

All data and code are open access and will be released at publication time.

## Acknowledgments

The research reported here was supported by AFOSR under grant #FA9550-16-1-0410, DARPA under contract #HR0011-19-9-0035 and NSF under grant #2024607.

## 5 Supplement

This section contains supplementary figures/tables with additional details in support of the material covered in the main text.

### Fig.2 Supplement

**Figure 11:**
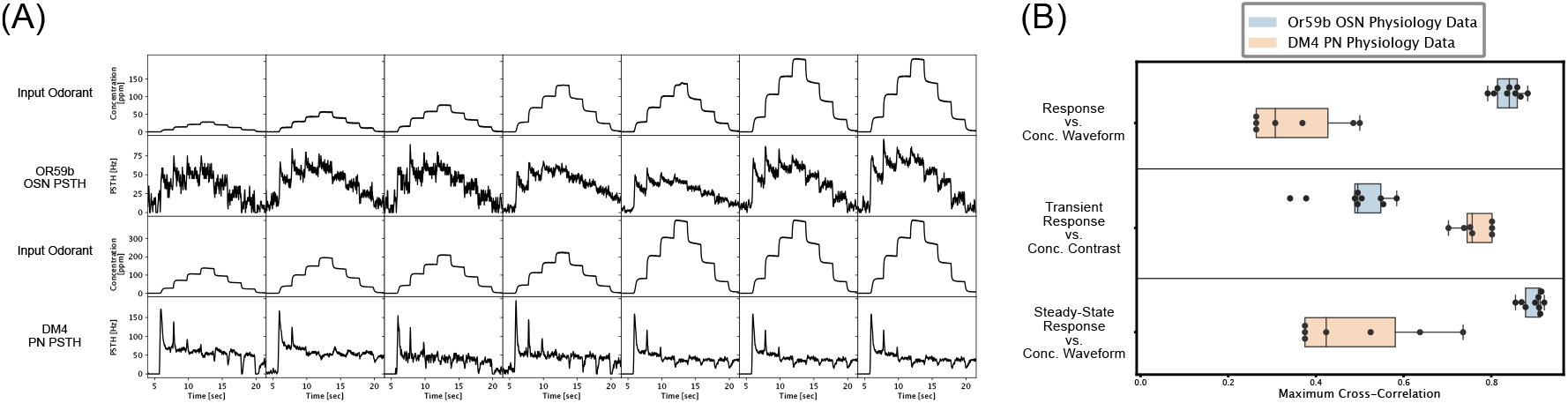
Concentration Invariance and ON/OFF Contrast Boosting for OR59b OSN and DM4 PN I/O pairs. (A) Input Acetone odorant waveforms and output PSTH for Or59b OSN and DM4 PN. Note that Or59b OSN and DM4 PN were recorded in different 7 sessions in [23]. The OSN recordings and PN recordings were matched for identical odorant waveform stimuli. (B) Maximum cross-correlation between different representations of odorant concentration waveforms (i.e., gradient, contrast) and different representations of neuron/model responses (i.e., transient, steady-state). Shown here is max*_τ_ ∫ x*(*t*)*y*(*t* + *τ*)*dt*, where *x*(*t*) is the steady-state/transient neuron response and *y*(*t*) is the concentration waveform/contrast.

### Fig.3 Supplements

**Figure 12:**
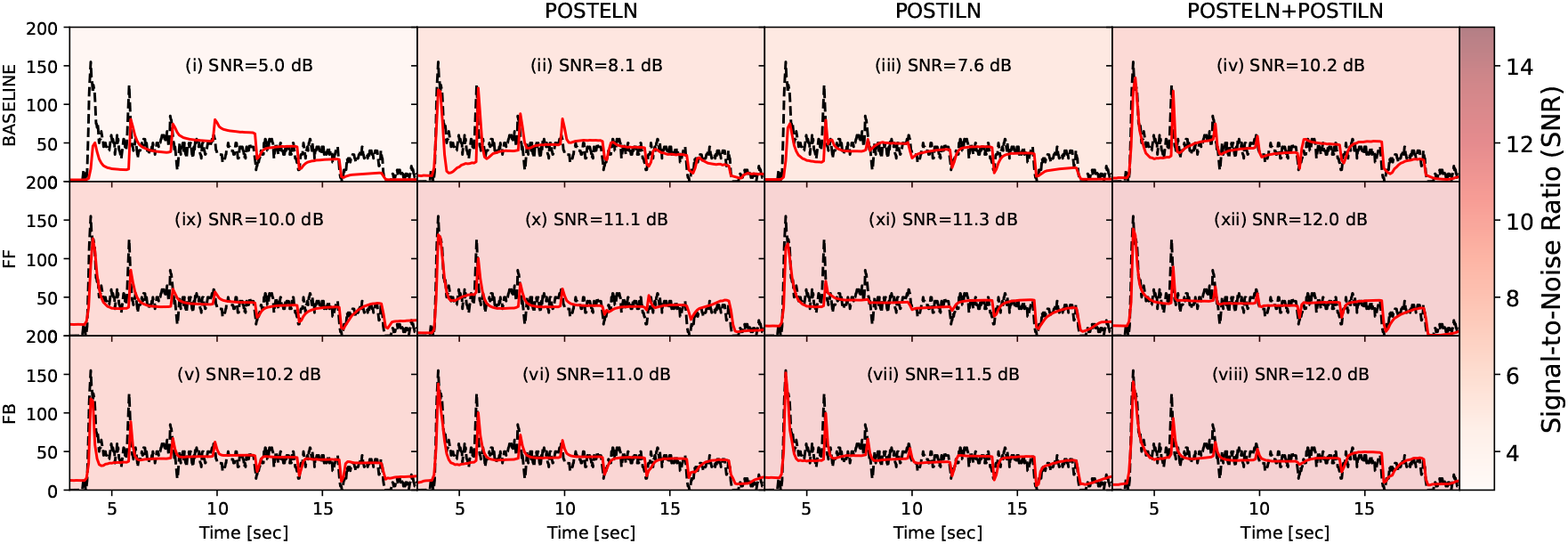
Best fitted temporal AL model responses to an Acetone staircase waveform. Each row represents a different model of Pre-LN Pathway (from top to bottom: Basline, Local-FF, Local-FB), and each column represents a different model of Post-LN Pathway (from left to right: No Post-LNs, Post-eLN only, Post-iLN only, Post-eLN + Post-iLN). Each sub-figure is colored by the overall SNR obtained by the model response (red line) vs. the physiology recording data (dashed black line).

**Figure 13:**
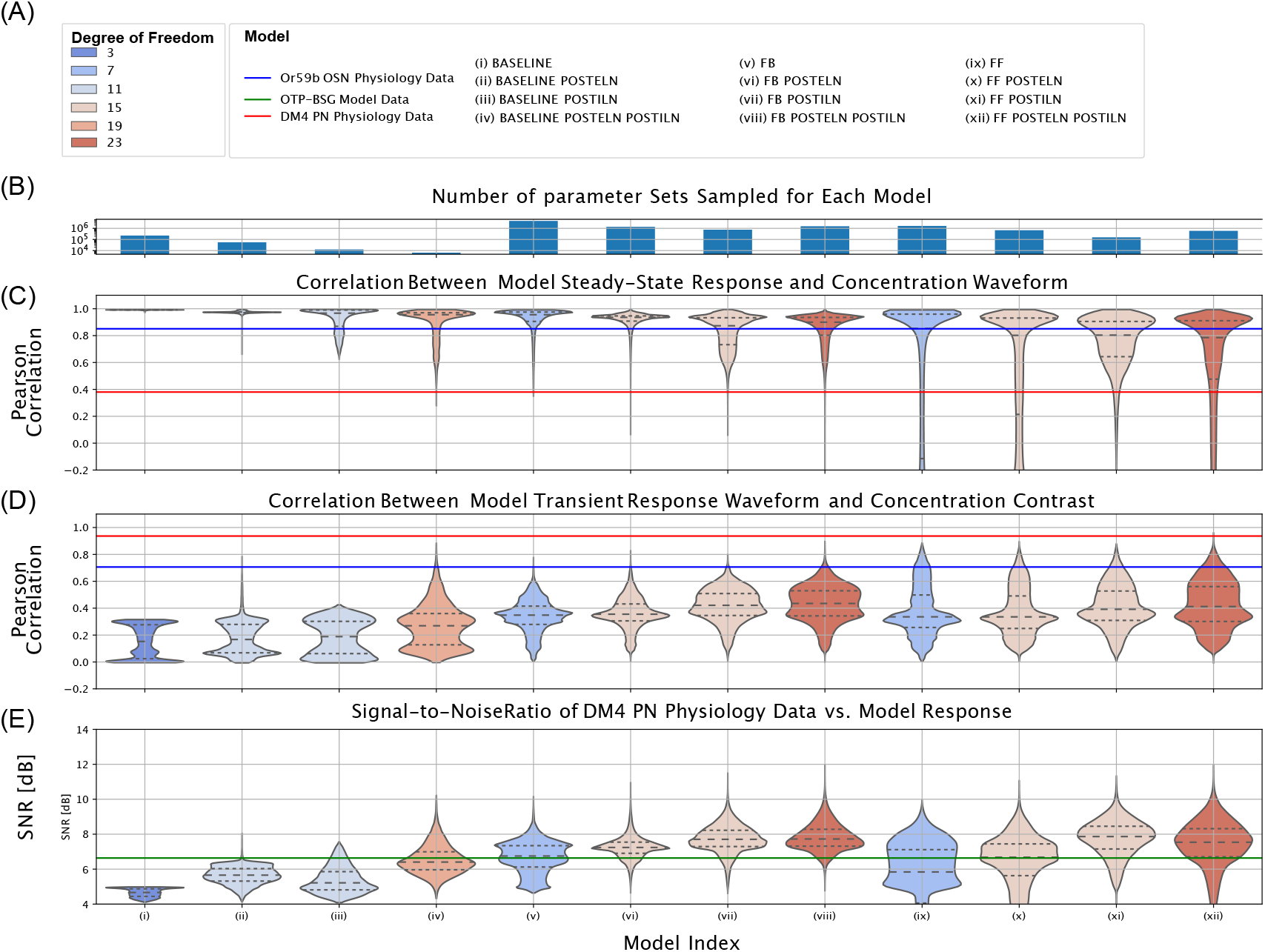
Comparison between 12 temporal processing circuits in the Antennal Lobe. (A) Legend for degree of freedom (color reference) and for circuit models. (B) Number of parameter sets sampled for each model; the y-axis is on the log-scale. (C) Sample distribution of the correlation between the steady-state responses and concentration waveform for each of the 8 models. The correlations between the steady-state response and the concentration waveform for DM4 PN and Or59b OSN physiology data are shown, for reference, in red and blue lines, respectively. (D) Sample distribution of correlation between the transient responses and concentration contrast for each of the 8 models. The correlations between the steady-state response and the concentration waveform for DM4 PN and Or59b OSN physiology data are shown, for reference, in red and blue lines, respectively. (E) Sample distribution of Signal-to-Noise Ratio between physiology recording data of DM4 PN and model responses. For reference, the Or59b OSN physiology data is linearly scaled to achieve the best SNR against the DM4 PN physiology data. The SNR between the scaled OSN response and PN response is shown in green.

### Fig.4 Supplements

**Figure 14:**
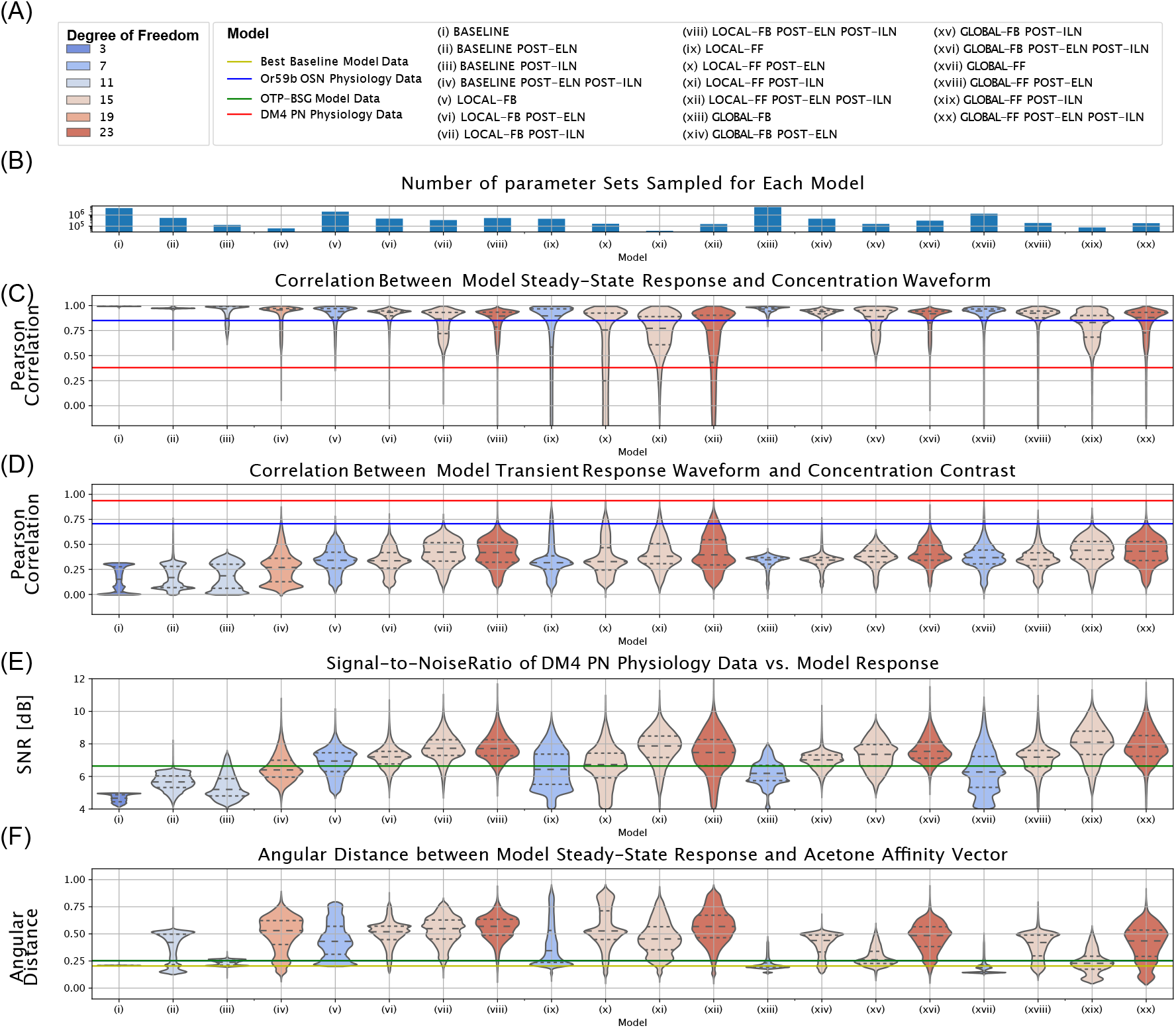
Comparison between all 20 spatio-temporal circuit models of AL. (A) Legend for model and degree of freedom. (B) Number of parameter sampled for each model; the y-axis is on the log-scale. (C) Correlation between the spatio-temporal model steady-state response of the DM4 PN to the staircase Acetone concentration waveform. (D) Correlation between the spatio-temporal model transient response of the DM4 PN to the staircase Acetone concentration contrast. (E) Signal-to-Noise ratio between spatio-temporal model response of the DM4 PN to the staircase Acetone concentration waveform and physiology data. The distance between the steady-state response of the OTP-BSG model and Acetone odorant affinity vector is shown in green for reference. (F) Average distance between spatio-temporal model responses and Acetone odorant affinity vector. The distance between steady-state response of the OTP-BSG model and the Acetone odorant affinity vector is shown, for reference, in green.

### Fig.9 Supplement

**Figure 15:**
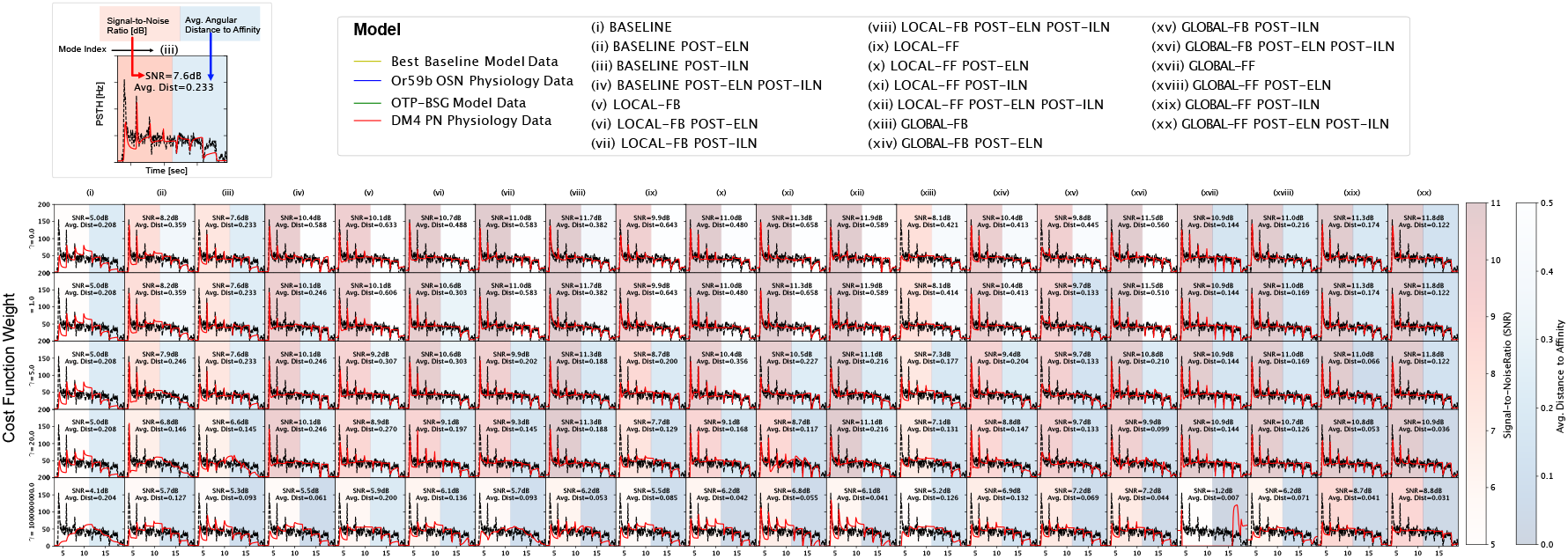
DM4 PN responses for all 20 spatio-temporal circuit models of the AL in **Fig**.14. Each row corresponds to a different weighting factor *γ* = {0, 1, 5, 20, 10^9^}. Each column shows the DM4 PN response of the best fit spatio-temporal model. (*top left*) Reference for each sub-figure. The background colors provide visual aids to compare signal-to-noise ratios (red) and average angular distances to affinity vectors (blue) across the circuit models and *γ* values.

**Figure 16:**
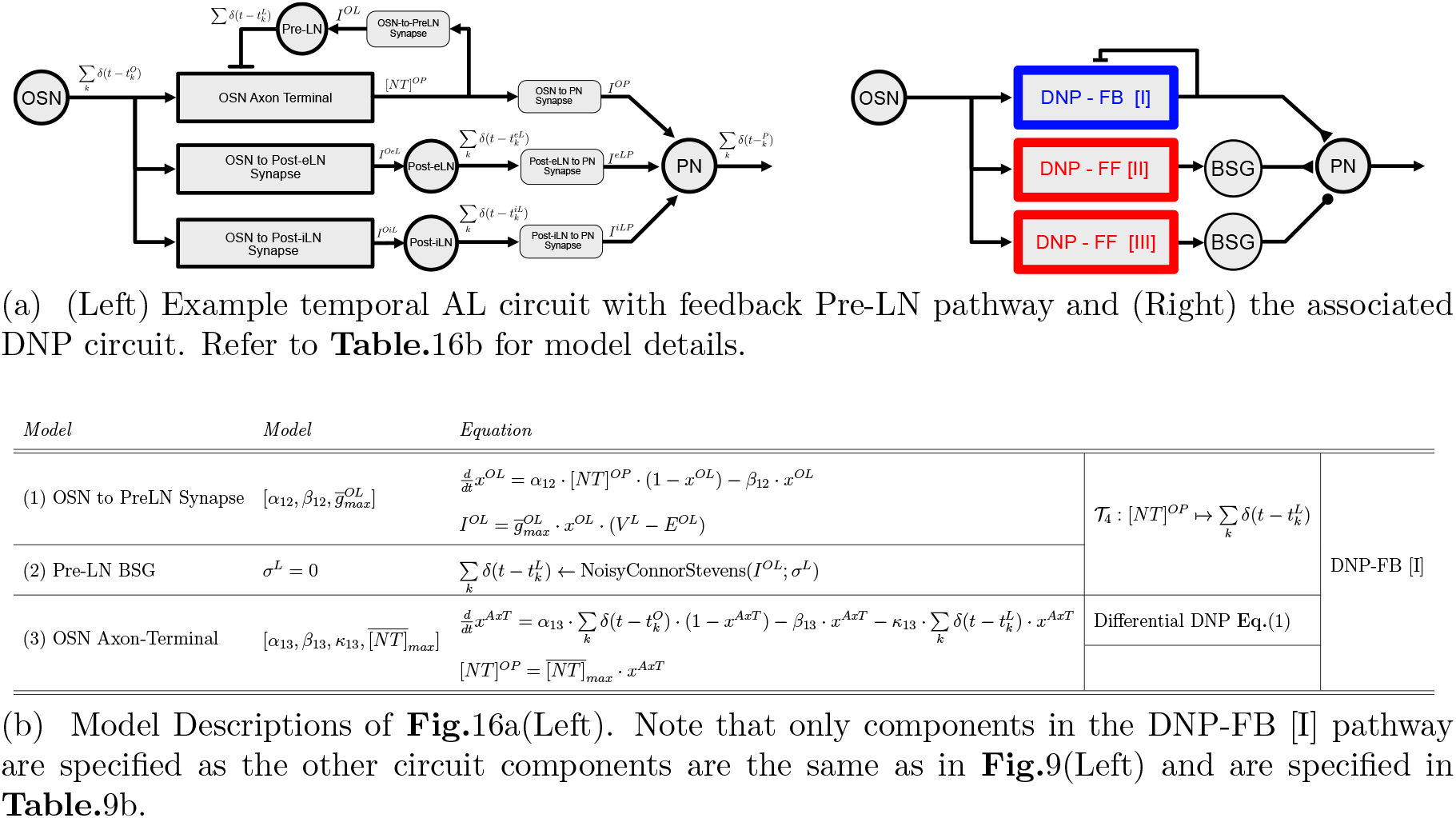
Example temporal AL circuit with feedback Pre-LN inhibition and associated equations.

### Fig.10 Supplement

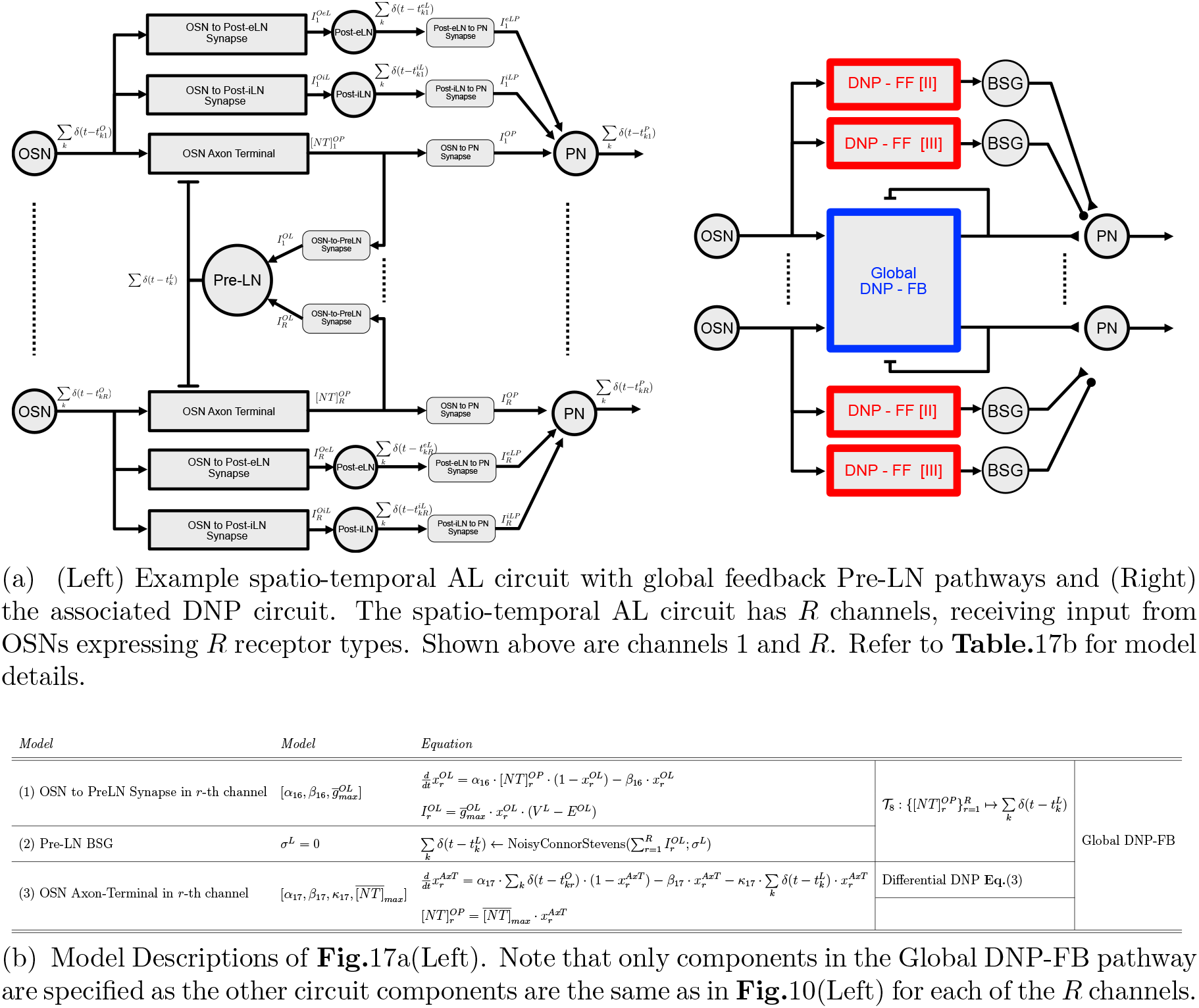

